# TRU-PE: A Universal, Trackable Prime Editor Toolkit for Robust Single-and Multi-Locus Genome Engineering

**DOI:** 10.64898/2026.02.20.706938

**Authors:** Zhichao Qiu, Keke Sun, Qingwei Zeng, Ziwei Luo, Xinran Liu, Yaping Li, Xiang Lei, Ruilin Zhao, Zhen Zhang, Damia Romero-Moya, Yuan Yang, Xuan Wang, Ehsan Hashemi, Joe Z. Zhang, Xiaolong Wang, Alessandra Giorgetti, Zhuobin Liang

**Affiliations:** Institute of Molecular Physiology, Shenzhen Bay Laboratory, Shenzhen, Guangdong, 518132, China; Regenerative Medicine Program, Bellvitge Institute for Biomedical Research (IDIBELL) and Program for Clinical Translation of Regenerative Medicine in Catalonia (P-CMRC), 08908, Barcelona, Spain; Department of Pathology and Experimental Therapeutics, Faculty of Medicine and Health Sciences, Barcelona University, Barcelona, 08907, Spain; Key Laboratory of Animal Genetics, Breeding and Reproduction of Shaanxi Province, College of Animal Science and Technology, Northwest A&F University, Yangling, Shaanxi, 712100, China; Department of Microbiology and Immunology, The Peter Doherty Institute for Infection and Immunity, University of Melbourne, Melbourne, Victoria, 3000, Australia; Department of Life Sciences and Biochemistry, Faculty of Arts and Science, Queen’s University, Kingston, Ontario K7L3N6, Canada; School of Science, Shenzhen Campus of Sun Yat-sen University, Shenzhen, Guangdong, 518107, China; Institute of Neurological and Psychiatric Disorders, Shenzhen Bay Laboratory, Shenzhen, Guangdong, 518132, China

## Abstract

Prime editing (PE) holds immense promise for precise genome editing, but its wider adoption is constrained by large payloads and inefficient delivery in recalcitrant cell types. Here, we present TRU-PE (Trackable, Robust, and Universal Prime Editor), a toolkit that overcomes these barriers via a split-vector architecture coupled with fluorescence-guided enrichment. This design ensures stoichiometric component expression, yielding marked improvements in editing efficiency and accelerated workflows compatible with both plasmid and viral delivery. By integrating a polycistronic hCtRNA-sgRNA/pegRNA array, the derivative TRU-MPE system enables simultaneous editing of up to ten genomic loci. Furthermore, it enhances the performance of evolved PE editors, allows flexible vector stoichiometry and inspires novel editing reporters. Leveraging these capabilities, we successfully generated a combinatorial human iPSC-based platform for GATA2 deficiency, which accurately recapitulates genotype-specific hematopoietic defects. TRU-PE thus provides a user-friendly, scalable framework for robust single- and multi-locus genome engineering, accelerating functional genomics and therapeutic development.

## Main

Prime editing (PE) has emerged as a transformative genome engineering technology, enabling the precise installation of targeted nucleotide substitutions, insertions, and deletions without requiring double-strand breaks (DSBs) or exogenous donor DNA templates^1^. This unique capability holds immense potential for fundamental research and the therapeutic correction of pathogenic mutations^2, 3^. However, while extensive protein and RNA engineering efforts have yielded substantial improvements in PE efficiency and accuracy^4^, bridging the gap between these enzymatic advancements and their practical application as robust, scalable tools for diverse genome editing tasks across different cellular contexts remains challenging.

A critical limitation restricting the broader adoption of PE is the inherent biophysical constraint of the editor payload combined with suboptimal enrichment strategies. The standard PE expression vector exceeds 12 kilobases (kb), which severely impedes plasmid delivery into biologically relevant but recalcitrant cell types, such as human induced pluripotent stem cells (hiPSCs), primary T cells, and other therapeutically significant lines^5–7^. Furthermore, conventional strategies rely on antibiotic resistance cassettes to select for cells successfully receiving the editor components. While simple and widely used, antibiotic selection has critical drawbacks for sensitive or precision editing: it is time-consuming, often suffers from high background due to non-expressing “escapees,” and can trigger cellular toxicity and p53 activation, which suppresses gene editing and compromises long-term viability^5, 8, 9^. These challenges highlight the need for a universal delivery architecture that can decouple editor performance from payload limitations.

Simultaneously, many severe medical conditions (*e.g.* cardiovascular diseases, myelodysplastic syndromes and neurodegenerative disorders), are driven by epistatic interactions between multiple genetic variants^10–12^. Interrogating these combinatorial networks requires the simultaneous installation of multiple specific mutations. However, efficient multi-locus PE in mammalian cells has remained a major hurdle. Existing multiplex PE approaches are typically restricted to modifying three or fewer loci simultaneously due to rapidly compounding biological and technical inefficiencies^13–15^. This constraint severely limits the field’s ability to model multifactorial diseases and orchestrate complex cellular engineering via PE.

To dismantle these barriers, we developed the Trackable, Robust, and Universal Prime Editor (TRU-PE) toolkit. Supported by a fully optimized and validated workflow, TRU-PE provides a user-friendly, easy-to-implement vector system that is broadly accessible to the scientific community through international repositories. By decoupling the bulky PE machinery into a size-balanced, tripartite architecture paired with distinct multi-fluorescent tracking, TRU-PE bypasses the bottlenecks of toxic antibiotic selection and payload delivery, enabling the precise, fluorescence-activated cell sorting (FACS)-based isolation of cells with optimal editor stoichiometry.

This optimal design yields context-specific enhancements in editing efficiency: up to 16-fold improvements in recalcitrant adherent lines (HeLa, MCF7) via plasmid transfection, up to 21-fold gains in sensitive pluripotent stem cells (hiPSCs), and 9- to 71-fold increases when adapting the system for lentiviral delivery in suspension T cells and HEK293T cells. Notably, it overcomes difficult loci and effectively unlocks substantial editing in traditionally intractable models. Crucially, the derivative TRU-MPE platform extends this capacity to enable the simultaneous editing of up to ten genomic loci. Furthermore, the TRU-PE architecture can act as a universal “operating system” for the rapidly evolving PE field, by seamlessly integrating with and maximizing the performance of state-of-art editors.

Finally, to demonstrate the platform’s capacity for complex disease modeling, we deployed TRU-MPE to dissect the multistage pathogenesis of GATA2 deficiency, a multisystem disorder where germline GATA2 heterozygous mutations cooperate with oncogenic somatic drivers causing myelodysplastic syndrome (MDS) and acute myeloid leukemia (AML). By simultaneously introducing a *GATA2* germline mutation alongside clinically recurrent secondary somatic mutations in *SETBP1* and *ASXL1*, we generated a comprehensive, combinatorial hiPSC-based cellular platform in a single editing round. Directed hematopoietic differentiation of these strictly isogenic models revealed clear genotype-specific phenotypic defects, suggesting a severe ASXL1-associated myeloid differentiation blockage.

## Results

### Optimization of vector architecture and selection strategies for robust prime editing

To establish a broadly applicable and highly efficient prime editing (PE) platform, we reasoned that the system must overcome three critical barriers: the toxicity and inefficiency of antibiotic selection, the delivery constraints of large plasmid payloads, and the requirement for scalable multiplexing. Consequently, we systematically evaluated three key parameters prior to finalizing our design: (1) strategies for the enrichment of edited cells, (2) validation of multiplexing-ready RNA polymerase III promoters, and (3) the impact of vector topology on delivery and editing outcomes.

We initiated our optimization in HEK293T cells by assessing the efficacy of puromycin-based selection on a conventional three-vector PE architecture (PE5max), incorporating GFP expression as an internal reporter for the delivery of nCas9 and reverse transcriptase (RT) (**Supplementary Fig. 1a**). We hypothesized that if puromycin selection is sufficiently robust, it would enrich for co-transfected cells that are GFP-positive. However, cells co-transfected with the three vectors using the recommended dosage ratio of 9:3:1 (nCas9-RT vs. pegRNA vs. nickingRNA-Puro), plateaued at approximately 80% GFP-positivity even with increasing puromycin concentrations (**Supplementary Fig. 1b,c**). To rule out co-transfection variability, we generated a unified vector (uPE5max-G) where the GFP and puromycin resistance markers were physically linked on the same backbone (**Supplementary Fig. 1a**). Theoretically, selection of this linked construct should yield 100% GFP-positive populations; however, the selected cells again capped at ∼80% GFP-positivity regardless of drug concentration (**Supplementary Fig. 1b,c**). Furthermore, high puromycin concentrations significantly reduced cell viability (**Supplementary Fig. 1c**).

These results indicate that puromycin selection is neither robust nor “clean,” suffering from both cytotoxicity and a high rate of non-expressing “escapees.” In contrast, flow cytometric analysis revealed that sorting for fluorescent marker expression consistently yielded populations with higher editing efficiencies compared to unsorted or antibiotic-selected controls (**Supplementary Fig. 1d**), implying that direct expression-based tracking is a more superior selection strategy to enrich for cells with higher editing potential.

Next, to establish a foundation for multiplexing, we investigated the feasibility of using the human cysteine tRNA (hCtRNA) promoter to drive expression of RNA elements, a prior strategy that enables polycistronic gRNA expression^14^. We constructed derivatives of the PE5max system where both the pegRNA and nicking sgRNA were driven by hCtRNA (tPE5max and tuPE5max) (**Supplementary Fig. 2a**). Editing assays confirmed that hCtRNA-driven systems achieved efficiencies matching those of conventional U6/H1-driven systems across multiple cell lines (**Supplementary Fig. 2b**,**c**), validating this promoter as a robust engine for subsequent multiplexing designs.

We next evaluated the impact of vector architecture on editing performance by comparing a unified “all-in-one” design against a split-vector configuration. To this end, we engineered fluorescently tagged variants of both the split (tdPE5max-GR; GFP and RFP) and unified (tuPE5max-G; GFP only) systems (**Supplementary Fig. 3a**). Although fluorescence-based enrichment significantly enhanced editing outcomes for both architectures relative to unsorted controls, achieving up to an 8.2-fold increase in HeLa cells (**Supplementary Fig. 3b,c**), the split-vector configuration consistently outperformed the unified design.

To elucidate the factors driving this difference, we examined the relationship between plasmid size and transfection efficiency. Transfection of cells with a panel of four GFP-tagged plasmids of increasing length revealed a marked inverse correlation between vector size and delivery efficiency, a trend that was particularly pronounced in recalcitrant cell lines such as HeLa and MCF7 (**Supplementary Fig. 4a,b**). These results indicate that while the unified vector provides design simplicity, its increased cargo size significantly impedes delivery; conversely, a split-vector architecture mitigates this barrier, thereby maximizing uptake efficiency in hard-to-transfect cell types.

### TRU-PE: a split-architecture, three-color tracking system for robust prime editing across cell types

Synthesizing the above findings, we established the TRU-PE (Trackable, Robust, and Universal Prime Editor) system. To strictly limit vector size below the ∼8–9-kb threshold identified as critical for efficient delivery (**Supplementary Fig. 4b**), TRU-PE adopts a tripartite split-architecture. The system redistributes nCas9, RT-MLH1dn, and the pegRNA-nickingRNA array across three size-balanced plasmids, coupling each to a distinct fluorescent reporter (BFP, GFP, and RFP) (**Fig. 1a**), to enable FACS-based isolation of cells with adequate PE component expression. Validation across 12 genomic loci revealed that this system achieves superior editing efficiency in diverse cellular contexts (**Fig. 1b–d**).

**Figure 1.**
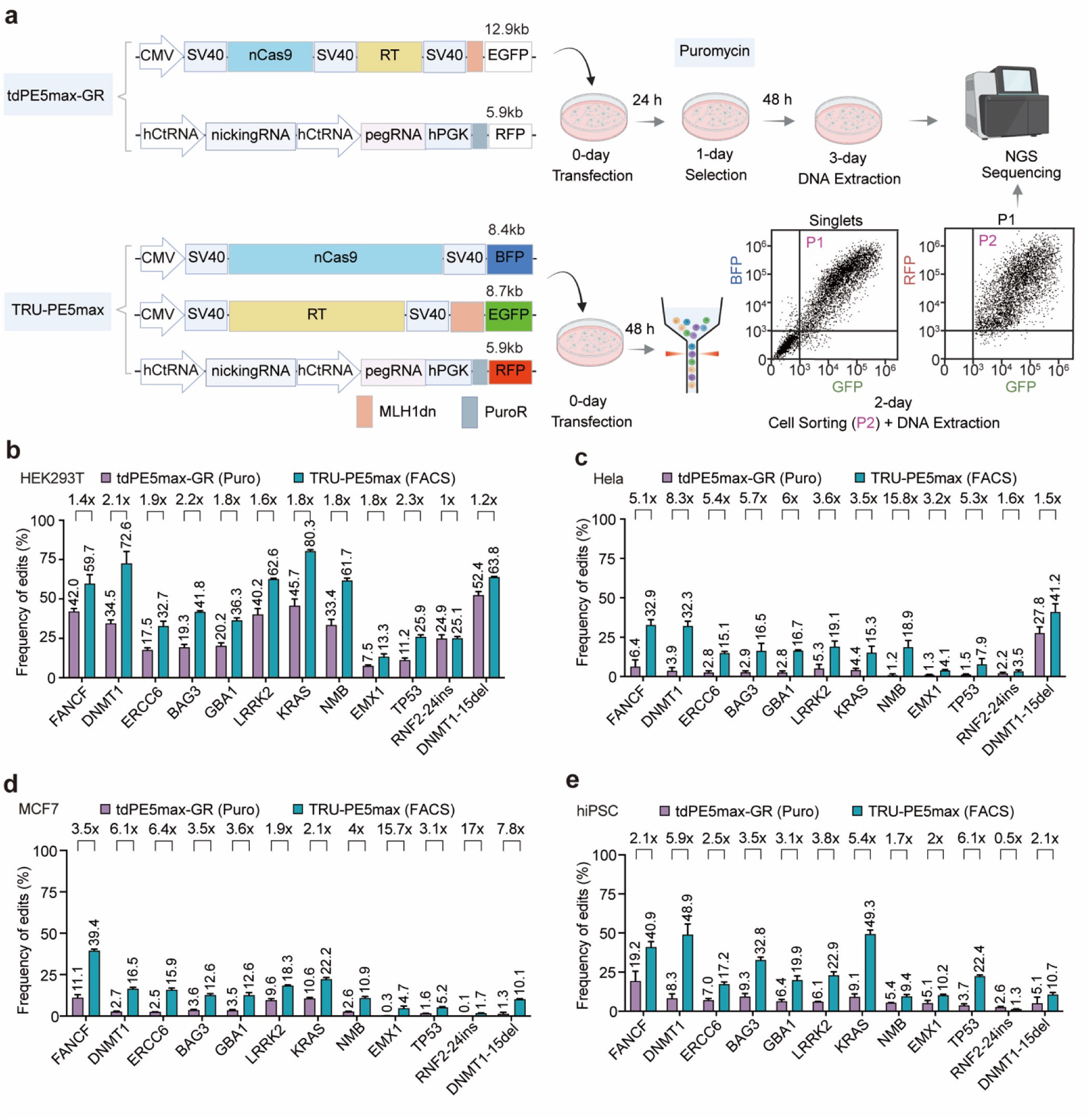
TRU-PE architecture enables robust prime editing via fluorescence-coupled component tracking. **a**, Schematics and workflows for control (tdPE5max-GR) and TRU-PE5max systems. Control cells are co-transfected with nCas9-RT and RNA plasmids (900 ng and 300 ng, respectively) followed by puromycin selection (24 h). TRU-PE5max splits components into nCas9 (BFP), RT (GFP), and RNA (RFP) vectors (400 ng each); triple-positive cells are enriched via FACS at 48 h (gating strategy P1 to P2). **b**–**e**, Comparison of editing efficiencies between puromycin-selected controls and FACS-enriched TRU-PE5max across 12 genomic loci in HEK293T (**b**), HeLa (**c**), MCF7 (**d**) cells and hiPSCs (**e**) . Puromycin was applied at 4 μg/mL for cancer lines and 0.5 μg/mL for hiPSCs. Fold-change improvements are indicated above bars. Editing outcomes were quantified via targeted deep sequencing. The associated unintended edit frequency is presented in **Supplementary Fig. 6**. Data are mean ± s.e.m. (n = 3 independent biological replicates).

While providing a modest ∼1.7-fold benefit on average in easily transfected HEK293T cells (**Fig. 1b**), TRU-PE proved transformative for recalcitrant lines. In HeLa cells, challenged by both intact DNA mismatch repair and limited transfectability, editing efficiencies increased by 1.5- to 15.8-fold compared to the puromycin-selected controls. Similarly, in transfection-refractory MCF7 cells, the system boosted editing rates by up to 15.7-fold (**Fig. 1c,d**), demonstrating that optimizing cargo topology is key to unlocking editing potential in difficult-to-transfect cells. To adapt TRU-PE to hiPSCs, we replaced the silencing-prone CMV promoter with EF1α promoter^16^ (**Supplementary Fig. 5a**). This modification restored robust component expression (**Supplementary Fig. 5b**) and translated into marked improvement (up to ∼6.1-fold) in editing efficiency compared to the standard PE controls (**Fig. 1e**). Importantly, this potency did not come at the cost of precision; deep sequencing confirmed that TRU-PE maintained a clean editing profile with minimal unintended edits (**Supplementary Fig. 6a–d**).

### TRU-MPE enables highly efficient simultaneous multi-locus editing

Next, we developed TRU-MPE for multiplex genome editing tasks by equipping the TRU-PE RNA vector with a polycistronic hCtRNA-sgRNA/pegRNA array (**Fig. 2a**). We validated this system in HEK293T cells by simultaneously targeting three and six distinct genomic loci, respectively, benchmarking performance against the previously reported Multiplex Prime Editing (MPE) system^14^ with the same hCtRNA-driven RNA array. Results showed that the TRU-PE architecture consistently outperformed the MPE controls, maintaining high efficiency even as the number of targets increased (**Fig. 2b; Supplementary Fig. 7**). Crucially, this multiplexing capability did not exhibit position-dependent biases (**Supplementary Fig. 8a,b**) or increased bystander mutation activity (**Supplementary Fig. 8a,b**).

**Figure 2.**
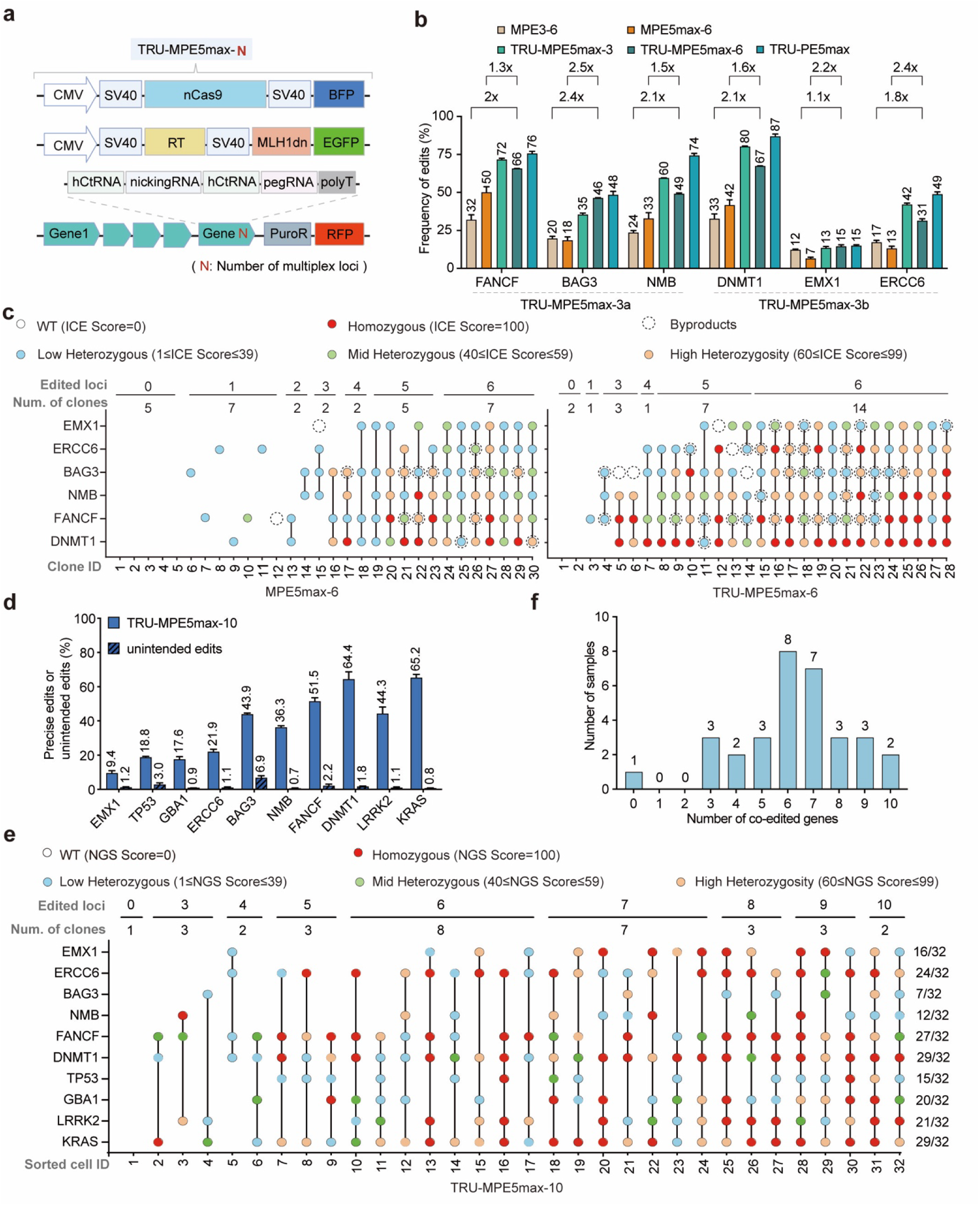
TRU-MPE facilitates scalable multiplex editing of up to ten genomic loci. **a**, Schematic of the TRU-MPE5max system. The RNA vector contains a polycistronic hCtRNA array driving the pegRNAs/sgRNAs. The red ‘N’ denotes the number of multiplexed targets. Components are co-transfected at a 1:1:1 ratio (400 ng each). **b**, Multiplex editing efficiencies of TRU-MPE at 3 and 6 loci in HEK293T cells compared to puromycin-selected MPE controls (MPE3 and MPE5max) and single-plex controls (TRU-PE5max). **c**, Clonal analysis of simultaneous 6-gene editing. Heatmaps display genotypes of individual clones expanded from single cells (MPE5max-6 vs. TRU-MPE5max-6). Genotypes are categorized by ICE scores. **d**, Simultaneous editing efficiencies at 10 distinct loci using TRU-MPE5max-10. **e**, Single-cell NGS analysis of 10-plex editing. Heatmap shows editing scores for sorted single cells (rows) across 10 loci (columns). Genotypes are categorized by NGS scores **f**, Frequency distribution of co-edited loci per cell from (**e**). Editing outcomes were quantified via targeted deep sequencing. The associated unintended edit frequency of **b** is presented in **Supplementary Fig. 8**. Data in **b**, **d** are mean ± s.e.m. (n = 3 independent biological replicates).

Clonal analysis confirmed the simultaneity of multiplex editing, demonstrating that TRU-MPE significantly outperformed the control MPE system by generating a greater proportion of clones with high editing completeness and homozygosity across all six targeted loci (**Fig. 2c**). Extending this capacity, we achieved successful simultaneous editing of up to ten genomic loci (**Fig. 2d**), and confirmed via a customized single-cell NGS pipeline (**Supplementary Fig. 9**) that approximately 6% of isolated cells (2/32) contained edits at all ten sites, while the majority harbored concurrent edits at 6 to 7 loci (**Fig. 2e,f**).

Recognizing that HEK293T cells often fail to recapitulate disease-relevant physiology^17,18^, we evaluated TRU-MPE in more biologically rigorous models. In HeLa cells and hiPSCs, the system achieved up to 7-fold and 21.1-fold increases in multiplexing editing efficiency, respectively, compared to the MPE controls (**Fig. 3a,b; Supplementary Fig. 10a,b**). Furthermore, the system supported robust multiplexing editing of ten loci in these recalcitrant lines without compromising editing purity (**Fig. 3c**; **Supplementary Fig. 10c**).

**Figure 3.**
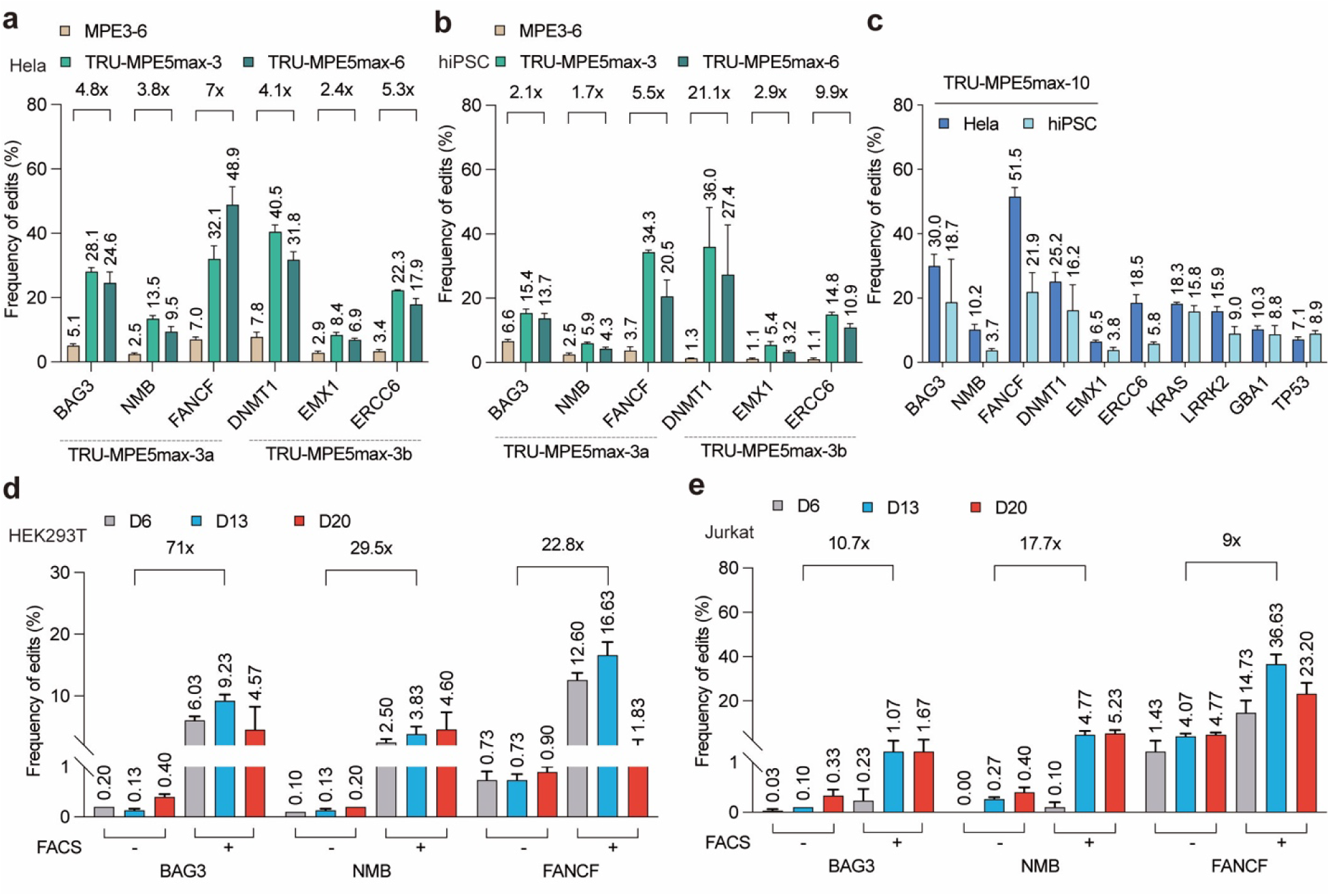
TRU-MPE enables high-efficiency multiplexing in recalcitrant cell types and supports viral delivery. **a**, **b**, Multiplex editing efficiencies in HeLa (**a**) and hiPSCs (**b**). TRU-MPE5max (3-plex and 6-plex) results are from FACS-enriched populations; MPE3-6 controls were puromycin selected. **c**, Simultaneous 10-plex editing efficiencies in HeLa and hiPSCs (TRU-MPE5max-10). **d**, **e**, Time-course analysis of lentiviral delivery for 3-plex editing (*BAG3*, *NMB*, *FANCF*) in HEK293T (**d**) and Jurkat T cells (**e**). Cells were transduced with MPE5max-3 or TRU-MPE5max-3 viruses and harvested at indicated days. Fold-changes at day 13 are shown. Editing outcomes were quantified via targeted deep sequencing. The associated unintended edit frequency is presented in **Supplementary Fig. 10**. Data are mean ± s.e.m. (n = 3 independent biological replicates).

Finally, to address delivery versatility, we packaged the TRU-MPE system into lentiviral vectors for the transduction of suspension cells (**Supplementary Fig. 11**). In Jurkat T cells and HEK293T cells, lentiviral TRU-MPE enhanced editing efficiencies by 9- to 17-fold and 22.8- to 71-fold, respectively, compared to the lentiviral MPE controls (**Fig. 3d,e**). Notably, substantial editing was readily achieved in the early days post-transduction, significantly shortening the experimental timeline compared to the 2-3 weeks typically required for conventional antibiotic selection workflows. Together, these findings validated TRU-MPE as a scalable and versatile toolkit for multiplex genome engineering.

### TRU-PE architecture is compatible with and enhances the performance of evolved prime editor variants

As the PE field advances, new variants with improved enzyme kinetics or structural stability continue to emerge. To ensure the TRU-PE platform remains a versatile toolkit for the community, we sought to demonstrate two critical features: first, that its split-architecture is broadly compatible with recent and future editor modifications; and second, that adapting these evolved editors into the TRU-PE format enhances their experimental workflow and editing performance compared to their original vector designs.

We first addressed the structural compatibility of the system using the recently described coiled-coil (CC) dimerization domains (P3/P4), which stabilize the split nCas9-RT interaction^19^. To assess structural compatibility, we engineered a TRU-PE5max-P3P4 variant (**Fig. 4a**), which maintained robust editing activity across multiple cell lines (**Fig. 4b–d**). Although the efficiency gains over the standard TRU-PE5max were modest compared to enhancements seen in conventional split-PE3 systems, likely due to the high RT stoichiometry in our fluorescence-enriched cells, these results confirm that the TRU architecture effectively accommodates P3/P4 dimerization domains without functional compromise.

**Figure 4.**
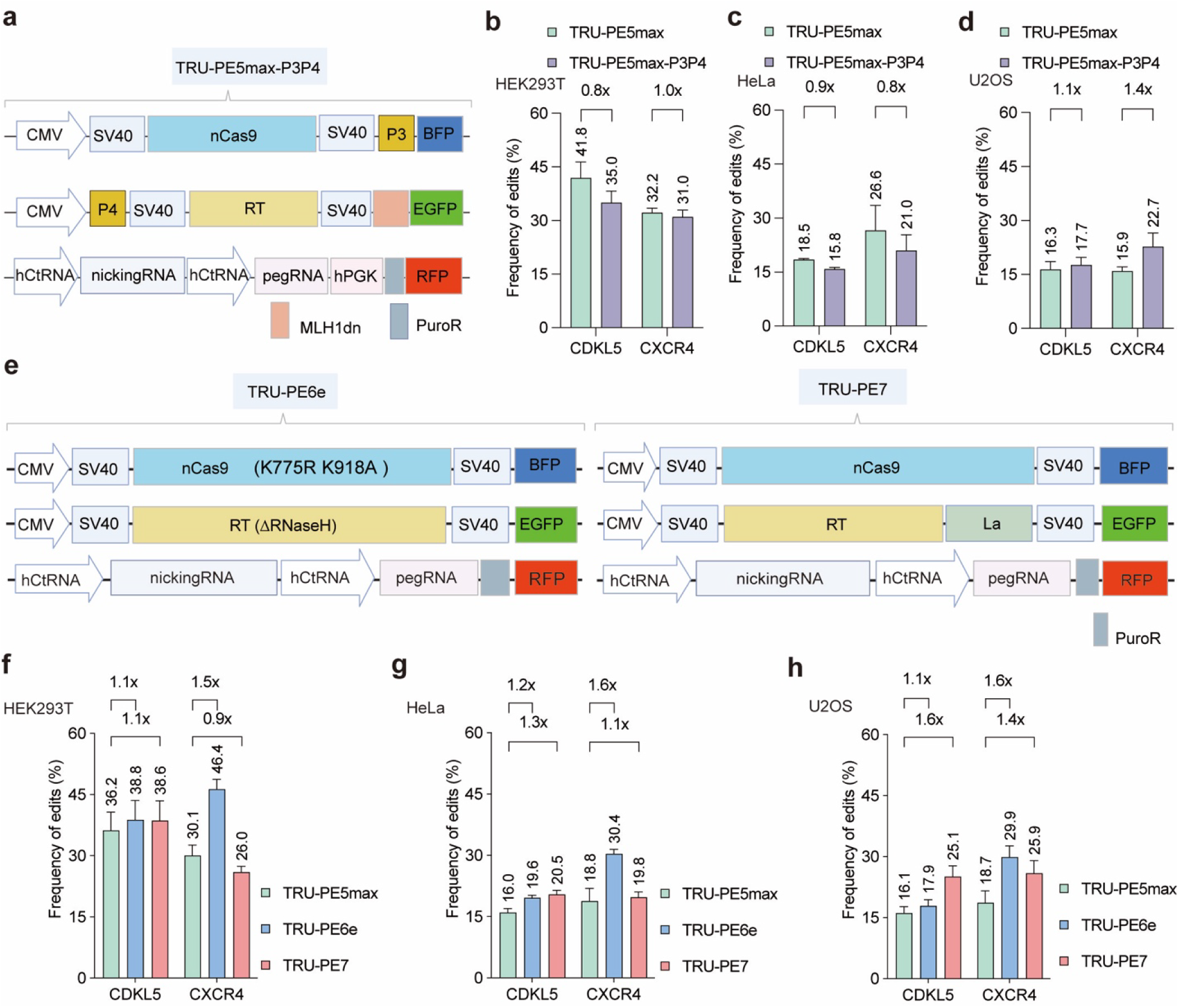
The TRU-PE architecture enhances the performance of evolved prime editor variants. **a**, Schematic of TRU-PE5max-P3P4 incorporating coiled-coil heterodimerization domains. **b**–**d**, Comparison of standard TRU-PE5max and TRU-PE5max-P3P4 efficiencies at *CDKL5* and *CXCR4* loci in HEK293T (**b**), HeLa (**c**), and U2OS (**d**) cells. **e**, Schematics of TRU-PE adapted for evolved editors PE6e (left) and PE7 (right). **f**–**h**, Editing efficiencies of TRU-PE5max, TRU-PE6e, and TRU-PE7 in HEK293T (**f**), HeLa (**g**), and U2OS (**h**) cells. All groups were FACS-enriched (triple-positive). Editing outcomes were quantified via targeted deep sequencing. Data are mean ± s.e.m. (n = 3 independent biological replicates).

Next, we investigated whether the TRU-PE architecture could actively enhance the performance of the latest generation of evolved editors: PE6e, a compact variant optimized through phage-assisted continuous evolution (PACE)^20^, and PE7, which integrates the La small RNA-binding protein^21^. We adapted these variants into the TRU-PE format (TRU-PE6e and TRU-PE7) and performed comparison against TRU-PE5max in HEK293T, HeLa, and U2OS cell lines (**Fig. 4e**). Strikingly, while the conventional PE6e and PE7 vectors exhibited limited editing efficiencies (approximately 5∼10%) at the selected challenging loci (*CXCR4* and *CDKL5*) as previously reported^20, 21^, likely due to the delivery limits of their full-length architecture and inefficient selection strategies, the TRU-PE adapted versions significantly outperformed them, yielding approximately a 5-fold improvement in editing efficiency at these sites (**Fig. 4f–h**). Therefore, by decoupling delivery from cargo size limitations and enabling precise stoichiometric enrichment, the TRU-PE architecture not only supports these advanced editors but maximizes their potential, offering a streamlined and high-performance workflow for utilizing state-of-the-art editing tools.

### Flexible vector stoichiometry and targeted reporter systems enhance prime editing precision

Conventional three-vector PE systems (*e.g.*, PE5max) typically necessitate a fixed 9:3:1 plasmid dosage ratio to accommodate major vector size discrepancies and ensure effective selection via the minimal puromycin-carrying vector. By uncoupling the components and tracking them individually, the TRU-PE architecture removes this constraint, providing the flexibility to customize vector stoichiometry for a convenient modulation of component expression levels in the cells. This is biologically advantageous: for example, while the RT enzyme requires high nuclear abundance to effectively drive reverse transcription, an excess of nCas9 can promote unwanted off-target nicking or the re-nicking of precisely repaired alleles. Similarly, an overabundance of pegRNA can trigger template over-amplification and error-prone repair, potentially leading to increased unintended edits in a sequence context-dependent manner ^22, 23^.

To test whether modulating vector ratios could improve editing purity, we evaluated a 6-loci multiplex editing panel in HEK293T cells while intentionally reducing the input of one or two vectors significantly below the standard 400 ng amount (**Fig. 5a**). Because the TRU-PE system tracks all components, we successfully enriched for triple-positive cells and achieved substantial editing even at suboptimal dosages. Crucially, as hypothesized, restricting the dosage of the nCas9 or pegRNA vectors, but not the RT-MLH1dn vector, significantly suppressed unintended edit frequencies at the BAG3 and FANCF loci, which are intrinsically prone to byproduct formation (**Fig. 5b**; **Fig. 2c**). Consequently, this customizable stoichiometry yielded a purer editing population characterized by substantially higher intended/unintended edit ratio (**Fig. 5c**).

**Figure 5.**
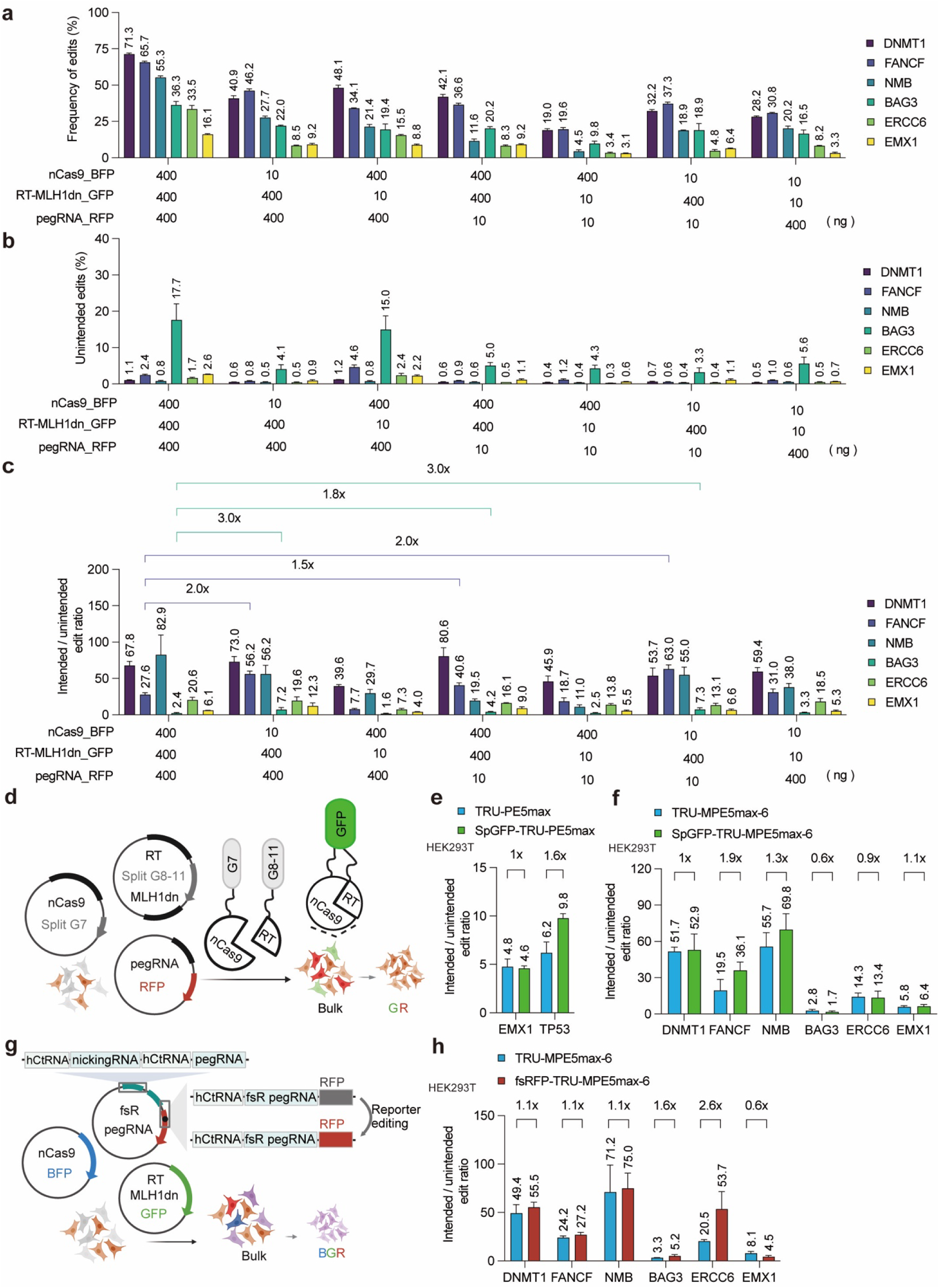
Tunable vector stoichiometry and fluorescent reporters optimize editing purity and enrichment. **a**–**c**, Effect of plasmid stoichiometry on editing outcomes in HEK293T cells. Editing efficiency (**a**), unintended edit frequency (**b**), and intended/unintended edit ratio (**c**) were measured under varying mass inputs (ng) of TRU-MPE5max-6 components. **d**, Schematic of SpGFP-TRU-PE using split-GFP reconstitution to track nCas9-RT assembly. **e**, **f**, Intended/unintended edit ratios using SpGFP-TRU-PE enrichment for singleplex (**e**) and 6-plex (**f**) editing. **g**, Schematic of fsRFP-TRU-PE, containing a frameshift RFP surrogate reporter on the pegRNA vector. **h**, Intended/unintended edit ratios using fsRFP reporter enrichment across 6 loci. Editing outcomes were quantified via targeted deep sequencing. Data are mean ± s.e.m. (n = 3 independent biological replicates) .

Building on this capacity to monitor component expression and tune stoichiometry, we next designed specialized fluorescent reporter systems to target and isolate cells that have successfully navigated critical rate-limiting steps of the PE process: editor assembly and functional editing. First, to capture cells with successful nCas9 and RT co-assembly in the nucleus, we developed a reporter system by fusing split-GFP fragments (GFP1-10 and GFP11) to the nCas9 and RT domains, respectively, to monitor the PE complex formation (**Fig. 5d**). Initial validations indicated a modest 1- to 2-fold improvement in editing efficiency in both single- or multi-locus editing (**Fig. 5e,f**).

Second, to directly isolate cells wherein the PE machinery is actively functional, we engineered a frameshift RFP (fsRFP) surrogate reporter into the pegRNA vector **(Fig. 5g**). In this design, successful editing of the surrogate target restores the RFP reading frame, linking fluorescence directly to functional PE activity. While baseline testing showed modest enhancements **(Fig. 5h**), we hypothesized that modulating the surrogate-pegRNA vector ratio could optimize the balance between target editing and surrogate repair. Together, these derivative TRU-PE designs demonstrate that the system’s architecture not only affords stoichiometric flexibility but also provides a versatile foundation for engineering advanced reporters that pinpoint distinct stages of the editing lifecycle.

### TRU-PE empowers hiPSC-based polygenic disease modeling through single-step combinatorial editing

To evaluate the utility of TRU-PE in establishing models for complex genetic disorders, we focused on GATA2 deficiency, a predisposing disorder that progresses to MDS and AML via the acquisition of secondary somatic mutations^24–26^. A recent study by our collaborative team detailed how specific secondary mutations, such as those in *SETBP1* and *ASXL1*, cooperate with germline *GATA2* mutations to alter leukemogenic trajectories^27^. While that study provided a comprehensive molecular characterization of these epistatic interactions, the underlying isogenic hiPSC models were generated using conventional homology-templated CRSIPR-Cas9 approach via iterative rounds of editing to install the multiple mutations. These sequential methods are labor-intensive, technically challenging in pluripotent stem cells, and prone to introducing culture-induced artifacts over prolonged passaging.

We hypothesized that the TRU-MPE platform could overcome these limitations by introducing all disease-relevant mutations in all possible combinations after a single editing round, thereby providing a robust starting point for deep mechanistic studies. We designed a multiplexed pegRNA array to concurrently install the primary germline mutation (*GATA2*^R396Q^) alongside two of the most recurrent secondary somatic mutations (*SETBP1*^D868N^ and the frameshift *ASXL1*^G646Wfs*12^)^28^ into a healthy donor derived hiPSC line (**Fig. 6a,b**). Remarkably, in a single round of editing and sorting, TRU-MPE5max successfully generated an extensive library of isogenic clones encompassing single, double, and triple mutants (**Fig. 6c**, **Supplementary Fig. 12**). Immunofluorescence analysis confirmed that all established lines maintained robust expression of key pluripotency markers (Nanog and SSEA4) comparable to wild-type controls, ensuring their suitability for downstream lineage specification (**Fig. 6d**). This simultaneous derivation approach significantly reduces the experimental timeline from months to weeks and providing a robust framework for engineering complex genetic interactions using hiPSCs and other stem cells.

**Figure 6.**
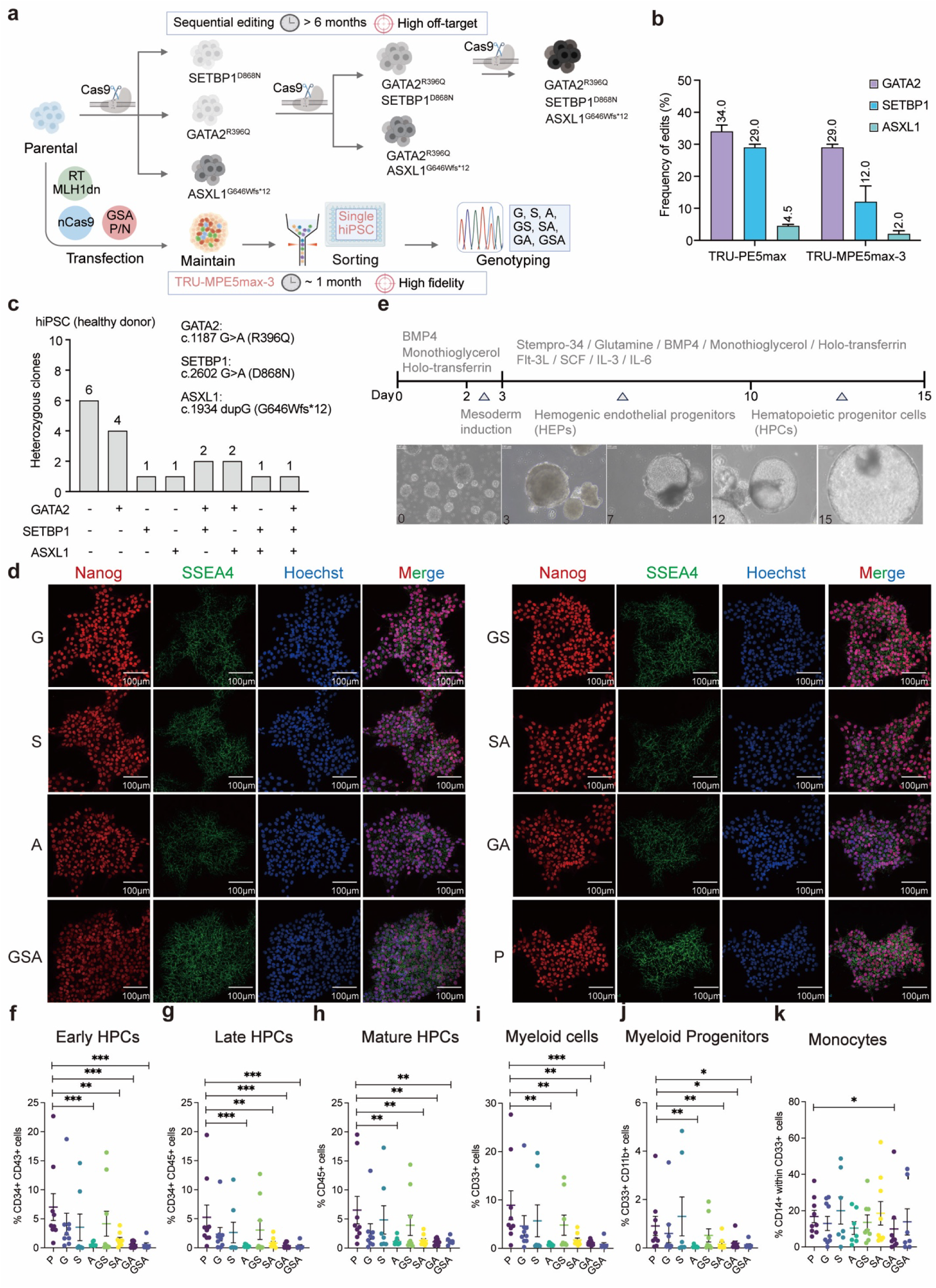
Single-step generation and functional validation of a combinatorial GATA2 deficiency hiPSC library for dissecting multistage hematopoietic pathogenesis. **a**, Comparison of conventional sequential editing (top) vs. single-step TRU-MPE5max strategy (bottom) for generating *GATA2, SETBP1,* and *ASXL1* mutants. **b**, Editing efficiencies at *GATA2, SETBP1,* and *ASXL1* loci using TRU-PE5max (single) or TRU-MPE5max (multiplex). **c**, Genotypic distribution of isolated hiPSC clones showing successful recovery of single, double, and triple mutants. **d**, Immunofluorescence of pluripotency markers (Nanog, SSEA4) in edited clones. Scale bars, 100 μm. **e**, The top panel outlines the differentiation timeline, specific developmental stages, and added cytokines. The bottom panel displays representative bright-field images of EBs at days 0, 3, 7, 12, and 15 of differentiation. **f**–**k**, Flow cytometric analysis of hematopoietic differentiation at day 15. Plots show frequencies of early HPCs (CD34^+^CD43^+^, **f**), late HPCs (CD34^+^CD45^+^, **g**), total leukocytes (CD45^+^, **h**), myeloid cells (CD33^+^, **i**), myeloid progenitors (CD33^+^CD11b^+^, **j**), and monocytes (CD14^+^, **k**). Genotypes: P (Parental), G (*GATA2*), S (*SETBP1*), A (*ASXL1*). Data are mean ± s.e.m (n = 7–9 independent biological replicates). Statistical significance was determined by two-sided Mann–Whitney test: (* p < 0.05, ** p < 0.01, *\*\*\** p < 0.001).

### Functional deconvolution of GATA2 deficiency phenotypes via TRU-PE generated isogenic hiPSC models

To functionally assess these TRU-PE generated isogenic hiPSC models and verify their consistency with prior iteratively edited lines, we subjected the isogenic library to directed hematopoietic differentiation, tracking the developmental progression from pluripotent cells to hematopoietic progenitors (**Fig. 6e**). The differentiation outcomes revealed a clear genotype-dependent hierarchy in hematopoietic output. In line with clinical GATA2 deficiency, GATA2 single mutants (G) displayed only a mild reduction in hematopoietic output compared to parental (P) controls. Conversely, multi-step clones (GS, GA, GSA) underwent a severe developmental blockade at early hematopoietic stages; specifically, the emergence of both early (CD34+CD43+) and late (CD34+CD45+) hematopoietic progenitor cells (HPCs) was markedly reduced (**Fig. 6f,g**). Consequently, the terminal yield of mature HPCs (total CD45+) was severely diminished specifically in these complex mutant backgrounds, rather than uniformly across all GATA2-deficient lines (**Fig. 6h**).

Beyond the primary GATA2 defect, the simultaneously generated library enabled a direct comparative analysis of the secondary mutations. The *SETBP1* single mutant (S) maintained robust hematopoietic potential, yielding early progenitor and mature populations at frequencies significantly higher than those observed in either the *ASXL1* (A) or *GATA2* (G) single mutants (**Fig. 6f–h**). In stark contrast, the *ASXL1* single mutant exhibited a severe developmental defect, restricting hematopoietic output to levels comparable to the GATA2-deficient lines. This impairment further extended into the myeloid lineage; the frequencies of total myeloid cells (CD33+), myeloid progenitors (CD11b+CD33+), and terminally differentiated monocytes (CD14+ within the CD33+ compartment) showed a pronounced downward trend across the mutant genotypes relative to the parental control, confirming severe disruptions in both early progenitor specification and downstream myeloid maturation (**Fig. 6i–k**). By efficiently recapitulating these complex, multi-lineage phenotypic trajectories in a strictly isogenic background, these data underscore the capacity of TRU-MPE to serve as a potent platform for accelerating the functional deconvolution of polygenic disease mechanisms and potential cell therapy engineering by correcting patient-derived stem cells.

## Discussion

The broad deployment of PE has been constrained by a delivery paradox: the editor’s versatility relies on a large, multi-component architecture that inherently restricts its delivery into biologically relevant, recalcitrant cell types. Various techniques to overcome this problem rely on specialized formulations or complex engineering, such as such as engineered viral-like particles (eVLPs)^29^, lipid nanoparticle (LNP)-encapsulated mRNA^30^, or purified ribonucleoprotein (RNP) complexes^31^. The TRU-PE system developed in this study provides a highly accessible and robust toolkit that overcomes these delivery and efficiency bottlenecks. By offering a straightforward, plasmid-based workflow compatible with standard laboratory settings, TRU-PE democratizes access to high-efficiency genome engineering for a broad user community, from basic researchers modeling disease to advanced users developing cell therapies.

To contextualize the advantages of TRU-PE, it is helpful to compare it with existing PE optimization strategies. Surrogate reporters, such as PEAR^32^ and fluoPEER^33^, successfully enrich for edited populations by providing a direct fluorescent readout of active editing. However, these trans-acting systems introduce an exogenous substrate that can kinetically compete with the endogenous target locus for editor binding. TRU-PE circumvents this competition by utilizing expression-coupled fluorescence, ensuring non-competitive enrichment. Furthermore, although previous split-PE architectures have successfully bypassed viral packaging limits^34^, their overall efficiencies are frequently compromised by the kinetic barriers of intracellular protein reconstitution. Our multi-channel FACS approach overcomes this by enriching for cells with oversaturated nuclear concentrations of the split components, thereby driving complex assembly through mass action.

To rigorously benchmark TRU-PE (**Fig. 1**), we utilized a fluorescently tagged, dual-vector system (tdPE5max-GR) as our baseline rather than the conventional non-fluorescent PE5max (a comparative analysis of which is provided in **Supplementary Fig. 3**). Because both the control and TRU-PE systems possess fluorescent reporters, this controlled design effectively isolated the variables of vector architecture and enrichment strategy (puromycin vs. FACS), preventing baseline differences from confounding the results. Furthermore, we strategically retained the puromycin resistance (PuroR) cassette on the TRU-PE pegRNA expression vector. This ensures cross-compatibility of the pegRNA vector between systems and preserves a traditional drug-selection route for laboratories lacking access to FACS equipment, or for applications where bulk selection is preferred, thereby broadening the platform’s utility.

Building on this robust foundation, the TRU-MPE system achieved high-efficiency simultaneous editing at up to ten loci (**Fig. 2** and **Fig. 3**), representing a substantial leap in scalability. While recent multiplexing PE advances, such as hCtRNA driven drive-and-process (DAP) CRISPR arrays^14^ and Csy4-dependent architectures^42^ represent major milestones, they frequently encounter a catalytic ‘dilution effect’ as targets increase or necessitate the co-delivery of exogenous processing RNases. TRU-MPE circumvents these bottlenecks through its decoupled architecture. By separating the components into size-balanced vectors, the system enriches for cells with an intracellular stoichiometric surplus of prime editing machinery. This abundance effectively buffers against multi-locus dilution, maintaining potent catalytic activity across multiple sites. Furthermore, to rigorously validate the editing simultaneity, we developed a customized single-cell NGS pipeline (**Supplementary Fig. 9**), which provides a low-cost, clonal expansion-independent method for rapidly profiling multi-locus editing events for FACS-harvested single cells. By bypassing the labor-intensive requirement of expanding single clones to obtain sufficient genomic DNA for target amplification, this pipeline not only confirms the TRU-MPE platform’s performance but also establishes a scalable, high-throughput standard for the field to benchmark multiplex editing tasks.

Such scalable multiplexing is critical for dissecting the ‘genome-phenome’ maps of complex polygenic traits. For example, it allows the simultaneous introduction of multiple GWAS-identified risk variants or modifier genes^36^, enabling the deconvolution of epistatic networks that are inaccessible to single-gene models. In the context of cell therapy, TRU-PE offers a streamlined solution for next-generation engineering. For instance, in CAR-T cell production, the system could enable the ‘one-shot’ insertion of CAR transgenes alongside the multiplex knockout of inhibitory receptors (*e.g.*, PD-1, CTLA-4) or endogenous TCRs (TRAC). This simultaneous genome insertion and alteration strategy is essential for generating potent, universal, and exhaustion-resistant cell products^33–35^. Besides biomedicine, this editing capacity lays the groundwork for synthetic biology applications, such as orchestrating genome-scale regulatory networks engineering^40^, executing synthetic genome debugging^37^, or constructing virtual cell models^38^.

Furthermore, we demonstrated that the TRU-PE architecture is compatible with and enhances evolved editor variants (**Fig. 4**). Interestingly, incorporating the P3/P4 heterodimerization domains into the TRU-PE architecture yielded only marginal improvements in editing efficiency (**Fig. 4b–d**), contrasting with previous reports emphasizing the necessity of physical tethering for split-PE assembly^19^. We attribute this not to structural incompatibility, but rather to fundamental differences in enrichment strategies. The multi-channel FACS enrichment employed by TRU-PE achieves exceptionally high local concentrations of nCas9 and RT proteins within the nuclei of sorted cells. Under such oversaturated stoichiometric conditions, the collision and assembly dynamics of the two proteins likely reach a plateau, rendering the marginal benefit of P3/P4-mediated physical tethering negligible. This observation indirectly highlights the inherent efficiency of the TRU-PE fluorescence-enrichment strategy in ensuring robust RNP assembly.

The TRU-PE system also demonstrates a unique capacity to control the formation of unwanted byproducts through simple vector stoichiometry (**Fig. 5**). Our data indicates that reducing the nCas9 vector input (*e.g.*, to 10 ng) while maintaining high pegRNA levels significantly increases the ratio of intended to unintended edits (**Fig. 5c**). Mechanistically, an excess of nCas9 could generate surplus DNA nicks; if the cellular machinery cannot repair these nicks promptly, the repair pathway defaults to bystander mutations (indels and mismatches). Therefore, the stoichiometric flexibility of TRU-PE allows users to titrate nCas9 levels downward, which sustains high editing efficiency while markedly improving the purity of the edited cell pool.

Finally, we leveraged TRU-MPE to generate an isogenic library of hiPSC disease models encompassing *GATA2* and its cooperating somatic mutations (*ASXL1*, *SETBP1*) in a single editing round (**Fig. 6**). Generating such complex multi-mutant lines has historically been a major bottleneck in iPSC disease modeling, typically requiring months of labor-intensive, sequential editing that risks genomic instability^43–45^.

Upon hematopoietic differentiation (**Fig. 6e**), GATA2-mutant lines exhibited compromised hematopoietic potential (**Fig. 6f–h**), consistent with patient pathology^46^. In agreement with recent findings by our collaborative team^27^, we observed that *SETBP1* mutations maintain a relatively robust hematopoietic output compared to isogenic control. However, we noted a discrepancy regarding the *ASXL1* mutation phenotype: while the previous study reported a milder impact, our data here demonstrate that the identical *ASXL1* frameshift mutation imposes a severe differentiation blockade comparable to GATA2 loss. This variance aligns with recent findings that the clonal dominance of *ASXL1* mutations is modulated by inherited variants disrupting a GATA-binding enhancer element, which tunes the MSI2 expression levels required for mutant expansion during clonal hematopoiesis^47^. Consequently, distinct germline genetic backgrounds of the different hiPSC lines used in our studies, rather than difference in genome editing strategies, likely underpin these phenotypic divergences.

While the current study primarily highlights the exceptional performance of TRU-PE in *ex vivo* mammalian cell engineering, the TRU-PE architecture offers transformative potential for germline editing and the generation of complex transgenic animal models. By leveraging its robust sorting capability in embryonic stem cells (ESCs) or donor fibroblasts, TRU-PE enables the isolation of highly pure, multi-gene edited populations prior to somatic cell nuclear transfer (SCNT) or embryo implantation^48, 49^. This workflow is particularly critical for engineering large gene-edited animals (*e.g.*, porcine, bovine and ovine species) to improve their agricultural traits or to create sophisticated surrogates that more accurately recapitulate human physiology and disease pathology than rodent counterparts^50^.

Furthermore, TRU-PE establishes a rigorous foundation for *ex vivo* cell therapy and functional genomics. In therapeutic contexts, the ability to precisely titrate editor stoichiometry and enrich for successful assembly is ideal for engineering human hematopoietic or mesenchymal stem cells (HSCs/MSCs) to treat hematological malignancies, metabolic disorders, or target aging-related pathways^51, 52^. Beyond stem cell transplantation, the platform facilitates the rapid construction of high-complexity, isogenic organoid biobanks. These models, derived from TRU-PE-edited hiPSCs, offer a powerful arena for high-throughput drug screening and toxicology testing, enabling the dissection of polygenic interactions in a human-relevant tissue microenvironment^53, 54^.

In conclusion, the TRU-PE system reported in this study represents a significant leap forward in genome engineering technology. By providing a versatile and highly accessible toolkit for the rapid generation of complex disease models and an enabling framework for diverse genome engineering applications, TRU-PE empowers researchers to tackle genomic challenges that were previously beyond reach, setting a new standard for precision and scalability in the field.

## Methods

### Mammalian Cell Culture

HEK293T, HeLa, MCF-7, and U2OS cells were cultured in Dulbecco’s Modified Eagle’s Medium (DMEM; Gibco, C11995500BT) supplemented with 10% (v/v) fetal bovine serum (FBS; TransGen Biotech, FS301-02) and 1× penicillin–streptomycin (Sangon Biotech, E607011-0100) in a humidified atmosphere containing 5% CO_2_ at 37 °C. For routine passaging, adherent cells were grown to ∼80–90% confluency, washed once with sterile Dulbecco’s phosphate-buffered saline (DPBS; Gibco, 14190250) without calcium and magnesium, and subsequently treated with 0.25% Trypsin-EDTA (Gibco, 25200072) for 3 min in the incubator. An equal volume of complete medium was added to terminate the digestion, and the cell suspension was then centrifuged at 200 × g for 3 min at room temperature. The cells were passaged onto a new dish at a 1:10 dilution and cultured, with medium changes every other day.

Jurkat cells were maintained in RPMI-1640 (Gibco, C11875500BT) supplemented with 10% FBS (TransGen Biotech, FS301-02) and 1× penicillin–streptomycin (Sangon Biotech, E607011-0100) and cultured in an incubator with 5% CO_2_ at 37 °C. To ensure logarithmic growth, the cell density was maintained between 1 × 10^5^ and 1 × 10^6^ viable cells/mL, and the cell concentration was not permitted to exceed 3 × 10^6^ cells/mL. For subculturing, the cell suspension was transferred to a sterile 15 mL conical tube and centrifuged at 150 × g for 5 min to pellet the cells. After centrifugation, the supernatant was discarded, and the cell pellet was gently resuspended in fresh, pre-warmed complete growth medium. The cell suspension was then diluted to a seeding density of 1 × 10^5^ viable cells/mL in new culture flasks.

The human iPSC line 022 (provided by Dr. Joe Z. Zhang laboratory) was cultivated in eTeSR (STEMCELL Technologies, 100-1215) using feeder-free culture protocols in 6-well plates (Corning, 3516) that were coated with growth factor-reduced Matrigel (Corning, 354277). To passage hiPSCs, the cells were washed twice with sterile DPBS (Gibco, 14190250) without calcium and magnesium, and then incubated with 0.5 mM EDTA (Beyotime, ST066) diluted in DPBS for 5 min at 37 °C in 5% CO_2_, followed by mechanical dissociation to obtain aggregates consisting of approximately 5–8 cells. The cells were then collected in 5 mL of cell culture medium, centrifuged at 200 × g for 3 min at room temperature, and resuspended in 2 mL of fresh culture medium. Cells were subsequently counted and seeded into 6-well plates at a density of 5 × 10^5^ viable cells per well in 2 mL of eTeSR medium supplemented with 10 μM Y-27632 dihydrochloride kinase inhibitor (MedChemExpress, HY-10583) to enhance cell survival. After 24 h, the medium was replaced with fresh eTeSR medium without Y-27632, and the medium was changed every 2 days thereafter.

### Plasmid Construction

Plasmids constructed in this study are detailed in **Supplementary Table 1**, and the associated primer sequences used for construction are listed in **Supplementary Table 2**. The pU6-pegRNA-GG-acceptor (LL036; Addgene 132777), pGL3-U6-sgRNA-PGK-puroR (LL037; Addgene 51133), pCMV-PE2 (LL001; Addgene 132775), pCMV-PEmax-P2A-hMLH1dn (LL003; Addgene 174828), pMD2.G (LL705; Addgene 12259), and psPAX2 (LL706; Addgene 12260) plasmids were obtained from Addgene.

### Molecular Cloning

To obtain the PCR fragment for plasmid construction, specific amplification was performed using the 2× Phanta Flash Master Mix (Dye Plus) (Nanjing Vazyme Biotech, P520) according to the manufacturer’s instructions. Briefly, a 50 μL PCR reaction system was set up, containing 25 μL of 2× Phanta Flash Master Mix, 2 μL of each forward and reverse primer (10 μM), 1 ng of the target plasmid as the template, and nuclease-free water to adjust the final volume. The PCR cycling conditions were as follows: initial denaturation at 98 °C for 30 sec; 35 cycles of denaturation at 98 °C for 10 sec, annealing at 60 °C for 5 sec, and extension at 72 °C for 5 s/kb; followed by a final extension at 72 °C for 1 min. PCR products from all reactions were identified via agarose gel electrophoresis and purified by gel extraction (Omega E.Z.N.A. Gel Extraction Kit, D2500-02) according to the manufacturer’s instructions.

For the double restriction enzyme digestion steps in plasmid construction, appropriate reaction buffers were selected using the NEBcloner online tool (https://nebcloner.neb.com/) to identify the optimal buffer for each enzyme pair, and the reaction was assembled according to the manufacturer’s instructions.

All T4 DNA ligation reactions for vector construction were performed under the following standardized conditions. A 20 μL reaction mixture was prepared on ice, containing 1 μL of T4 DNA Ligase (New England Biolabs, M0202L), 2 μL of 10× T4 DNA Ligase Buffer (New England Biolabs, B0202S), 0.02 pmol of vector DNA, and 0.06 pmol of insert DNA. Nuclease-free water was added to bring the total volume to 20 μL. The reaction was gently mixed by pipetting, briefly centrifuged, and then incubated at room temperature for 1 h. Following incubation, the ligase was heat-inactivated at 65 °C for 10 min. Finally, the mixture was chilled on ice and used for transformation.

Gibson assembly was carried out with the NEBuilder HiFi DNA Assembly Master Mix (New England Biolabs, E2621L) to construct the different plasmid vectors by assembling the vector backbone with insert fragments, following the manufacturer’s instructions. Briefly, the Gibson assembly reaction was set up on ice according to the number of DNA fragments being assembled. For 2–3 fragments, a total of 0.1 pmol of the mixed DNA fragments was combined with 5 μL of the DNA assembly master mix. Nuclease-free water was added to bring the total reaction volume to 10 μL. The reaction was then incubated in a thermocycler at 50 °C for 15 min. For 4–6 fragments, a total of 0.3 pmol of the mixed DNA fragments was used with 5 μL of the Master Mix, and the volume was similarly adjusted to 10 μL with nuclease-free water. The incubation time at 50 °C was extended to 60 min. Following the incubation, all samples were placed on ice for subsequent transformation.

In the TRU-PE system, all plasmids constructed using pGL3-(2 BsaI)-PGK-puroR-mRFP1 (LL470) as the backbone vector were generated using the Golden Gate assembly method. Specifically, a 20 μL Golden Gate reaction mixture was prepared, containing the purified DNA fragments, 2 μL of 10× T4 DNA Ligase Buffer (New England Biolabs, B0202S), 2 μL of 10× rCutSmart Buffer (New England Biolabs, B6004S), 1 μL of T4 DNA Ligase (New England Biolabs, M0202L), and 1 μL of BsaI-HFv2 (New England Biolabs, R3733L). For assemblies involving 2 to 5 inserts, the mixture was subjected to 30 temperature cycles of 37 °C for 1 min and 16 °C for 1 min, followed by a final digestion step at 60 °C for 5 min.

When assembling 6 to 10 inserts, the protocol was adjusted to 30 cycles of 37 °C for 5 min and 16 °C for 5 min, concluding with the same final incubation at 60 °C for 5 min. Following the incubation, all samples were placed on ice for subsequent transformation.

### Transfection

HEK293T, HeLa, MCF7, and U2OS cell lines were cultured and transfected using linear polyethylenimine (PEI, MW 40,000, Yeasen, 40816ES02). One day prior to transfection, cells were seeded into 24-well plates to achieve 60–70% confluency at the time of transfection. Transfection mixtures were prepared by diluting plasmid DNA in 50 μL of Opti-MEM I Reduced Serum Medium (Gibco, 31985062) per well.

The specific DNA composition varied depending on the gene-editing system employed. For the standard PEmax system, each well received 900 ng of the prime editor plasmid, 300 ng of the pegRNA plasmid, and 100 ng of the nicking-gRNA plasmid. In the TRU-PE system, the transfection mixture comprised 400 ng of the nCas9-BFP plasmid, 400 ng of the RT-MLH1dn plasmid, and 400 ng of the hCtRNA- sgRNA/pegRNA plasmid. For other two-plasmid gene-editing systems, the total DNA amount consisted of 900 ng of the prime editor plasmid and 300 ng of the hCtRNA-sgRNA/pegRNA plasmid. Finally, for single-plasmid gene-editing systems, 900 ng of the respective plasmid was used. Following DNA preparation, PEI (1 mg/mL stock solution) was added to the diluted DNA at a DNA-to-PEI mass ratio of 1:2. The DNA–PEI mixture was incubated at room temperature for 15 min to allow complex formation before being added dropwise to the cell culture supernatant.

Human iPSCs were transfected using Lipofectamine Stem Transfection Reagent (Invitrogen, STEM00003) according to the manufacturer’s protocol. Briefly, plasmid DNA and 2 μL of Lipofectamine Stem Reagent were diluted separately in 25 μL of Opti-MEM I Medium per well. The diluted DNA was then combined with the diluted reagent and incubated for 15 min at room temperature.

Prior to adding the transfection complex, the culture medium was aspirated and replaced with 0.5 mL of Opti-MEM I Medium supplemented with CloneR2 (STEMCELL Technologies, 100-0691) to support cell survival. After a 4 h incubation at 37 °C with 5% CO_2_, 0.5 mL of room temperature eTeSR medium was added to each well. The cells were returned to the incubator overnight. The following day, the medium was replaced with 0.5 mL of fresh eTeSR medium. At 48 h post-transfection, cells expressing fluorescent proteins were sorted by FACS.

### Lentivirus Packing

To produce lentiviral particles, HEK293T cells were seeded at a density of approximately 1.2 × 10^7^ cells per 15 cm dish in 20 mL of complete medium. After 16 h, the cells were transiently transfected with a plasmid mixture containing 22.5 μg of the transfer plasmid, 19.1 μg of the packaging plasmid psPAX2, and 5.6 μg of the envelope plasmid pMD2.G. Transfection was carried out using linear PEI 40,000 at a working concentration of 1 mg/mL, in accordance with the manufacturer’s instructions. The culture medium was replaced with fresh complete medium 6 h post-transfection. Viral supernatants were harvested at 72 h and clarified by centrifugation at 400 × g for 10 min at 4 °C to remove cellular debris, followed by filtration through a 0.45 μm PES filter (Biosharp, BP-PES-45). Subsequently, viral particles were pelleted by centrifugation at 15,000 × g for 2 h at 4 °C. The resulting viral pellets from each dish were resuspended in 200 μL of RPMI-1640 medium, aliquoted into 50 μL volumes in 1.5 mL centrifuge tubes, and stored at –80 °C until use.

### Fluorescence-activated Cell Sorting (FACS)

To enrich for transfected cells, FACS was performed at 48 h post-transfection. The culture medium was carefully aspirated from each well, and cells were dissociated by adding 300 μL of trypsin and incubating at 37 °C for 2 min. Digestion was terminated by the addition of 500 μL of complete culture medium, and the mixture was gently pipetted to generate a single-cell suspension. The cells were then collected and resuspended in 200 μL of Flow Cytometry Staining Buffer (eBioscience, 00-4222-57) and maintained on ice. Prior to analysis, the cell suspensions were filtered through Falcon 5 mL polystyrene round-bottom tubes equipped with cell-strainer caps (Corning, 352235) to ensure a uniform single-cell suspension. Fluorescence intensity was assessed using a CytoFLEX SRT flow cytometer (Beckman Coulter) in the FITC-A, ECD-A, and PB450-A channels.

For the analysis of hematopoietic differentiation on day 15, embryoid bodies (EBs) were dissociated into single-cell suspensions using a two-step enzymatic digestion protocol. First, EBs were transferred to a 15 mL conical tube. The culture medium was aspirated, and 1.5 mL of Collagenase B (Sigma, 11088807001, final concentration 0.4 U/mL) was added. The mixture was incubated at 37 °C for 1 h. After removing the supernatant, EBs were incubated with 1 mL of 0.25 mM EDTA for 15 min at 37 °C, followed by centrifugation at 1,200 rpm for 5 min. The cell pellet was resuspended in FACS buffer, mechanically disrupted by pipetting, and filtered through a cell strainer.

Cells were stained with the following fluorophore-conjugated antibodies: CD34-PE (Becton Dickinson, 550761), CD43-APC (Becton Dickinson, 560198), CD45-BV421 (Becton Dickinson, 563879), CD11b-PE (Becton Dickinson, 555388), CD14-APC (Becton Dickinson, 555399), and CD33 (ABclonal, A22639). Cell viability was assessed using the Zombie NIR Fixable Viability Kit (BioLegend, 423105) prior to analysis on the flow cytometer.

### Genomic DNA Extraction

Approximately 6,000 sorted cells were collected directly into 30 μL of QuickExtract DNA Extraction Solution (Lucigen, QE09050). The samples were lysed by incubation at 65 °C for 6 min, followed by an inactivation step at 98 °C for 2 min. The resulting lysate was transferred to a new tube and used for subsequent PCR amplification and sequencing analysis.

### Analysis of Gene Editing via Sanger Sequencing Analysis

Target genomic loci were PCR-amplified from extracted genomic DNA using locus-specific primers. Amplicons were purified and Sanger sequenced. To quantify editing efficiencies and determine genotypes, sequencing chromatograms (.ab1 files) were analyzed using the Inference of CRISPR Edits (ICE) tool (EditCo; https://ice.editco.bio/). Experimental traces were aligned against unedited wild-type controls. For single-cell clonal analysis, individual clones were assigned to genotypic categories (e.g., wild-type, low/medium/high heterozygous, or homozygous) based on their computed ICE scores and sequence contributions.

### Analysis of Gene Editing via Amplicon Deep Sequencing

Deep sequencing reads were demultiplexed using fastq-multx with the barcode mismatch threshold set to 0. Following demultiplexing, only samples yielding more than 20,000 total reads were retained for subsequent analysis. For quality control, reads were filtered to ensure a minimum average quality score (-g) of 30 and a minimum single-base quality score (-s) of 10. Amplicon sequences (listed in **Supplementary Table 3)** were aligned to the reference sequence using CRISPResso2 in prime editing mode. The minimum homology for alignment was set to >60%^55^. Only samples exhibiting an alignment rate (aligned reads divided by quality control reads) of ≥90% were considered reliable and included in the final dataset. For the quantification of single-base substitutions, 25 bp were excluded from both the 5′ and 3′ ends of the amplicon sequence to minimize edge effects. Input parameters included the pegRNA spacer, pegRNA extension (containing the RT template with the edit and the primer-binding site), pegRNA scaffold, and nicking sgRNA sequences. The quantification window was defined to cover 5 bp centered around the 3′ end of the pegRNA extension (totaling 10 bp), along with an additional 2 nt flanking the cut sites of both the pegRNA spacer and the nicking sgRNA. For insertion prime edits, CRISPResso2 was executed using the same parameters, but with the intended edited sequence provided as the prime-edited reference sequence (--prime_editing_override_prime_edited_ref_seq). Prime editing efficiency was calculated as the percentage of reads containing the intended edit without unintended edits relative to the total number of reads aligned to the amplicon. Unintended edit frequency was quantified as the ratio of unintended edit-containing reads to the total number of reference-aligned reads. The PCR primers used for sequencing library preparation are listed in **Supplementary Table 4**.

### Targeted Multi-loci Amplification from Single Cells

To evaluate multiplexed editing efficiencies across ten distinct genomic loci, HEK293T cells were co-transfected with a triple-plasmid system (plasmids LL392, LL622, and LL1450). At 48 h post-transfection, single cells were sorted by FACS into 96-well PCR plates containing 20 μL of QuickExtract DNA Extraction Solution per well. Eight wells containing only lysis buffer were reserved as negative controls. Genomic DNA extraction was performed in a thermal cycler using the following program: 65 °C for 6 min, followed by enzyme inactivation at 98 °C for 2 min.

A first-round PCR master mix was prepared to simultaneously amplify the ten target gene loci. Each 50 μL reaction consisted of 25 μL of 2× Rapid Taq Master Mix (Vazyme, P222-01), nuclease-free water, and 0.2 μL of each primer pair targeting BAG3 (OL5527/5528), ERCC6 (OL4900/4901), EMX1 (OL5640/5641), NMB (OL5531/5532), DNMT1 (OL5533/5534), FANCF (OL5002/5003), GBA1 (OL5489/5490), LRRK2 (OL5491/5492), KRAS (OL5493/5494), and TP53 (OL5495/5496). Thermal cycling conditions were set as follows: initial denaturation at 95 °C for 5 min, followed by 40 cycles of 95 °C for 10 sec, 55 °C for 20 sec, and 72 °C for 15 sec, with a final extension at 72 °C for 5 min.

The first-round PCR products served as templates for the subsequent library construction steps. A 10 μL aliquot from each first-round reaction was distributed into ten separate second-round PCR reactions to individually amplify each locus using linker-containing primers. These primers included OL5788/5235 for BAG3, OL5789/5239 for ERCC6, OL5790/5611 for EMX1, OL5038/5791 for NMB, OL5535/5792 for DNMT1, OL5793/5043 for FANCF, OL5794/5498 for GBA1, OL5499/5795 for LRRK2, OL5501/5796 for KRAS, and OL5797/5798 for TP53. Each 10 μL reaction contained 0.2 μL of the specific primer pair and 5 μL of 2× Rapid Taq Master Mix, with thermocycling conditions identical to those used in the first round.

Subsequently, a 1 μL aliquot of the second-round product was used as the template for the third-round PCR. Barcoded primers (forward: OL3756–OL3851; reverse: OL4047–OL4142) were employed to enable sample multiplexing. Each 25 μL reaction consisted of 0.5 μL of each primer and 12.5 μL of 2× Phanta Flash Master Mix (Vazyme, P520). The amplification protocol comprised an initial denaturation at 98 °C for 3 min, 7 cycles of 98 °C for 10 sec, 60 °C for 10 sec, and 72 °C for 10 sec, followed by 8 cycles of 95 °C for 10 sec, 60 °C for 10 sec, and 72 °C for 10 sec, and a final extension at 72 °C for 5 min. Final PCR amplicons were pooled in volumes proportional to band intensities observed on agarose gels. The resulting library was subjected to next-generation sequencing, and data were analyzed as described in the previous section. The sequences of all PCR primers used in this study are listed in **Supplementary Table 5**.

### Immunofluorescence Staining

To assess the pluripotency of the edited hiPSC lines, cells were seeded onto Matrigel-coated 24-well glass-bottom plates. Upon reaching approximately 50% confluency, cells were fixed with 4% paraformaldehyde (PFA; Beyotime, P0099) for 15 min at room temperature, followed by permeabilization with 0.2% Triton X-100 (Solarbio, T8200) for 30 min at room temperature. Cells were then blocked with 5% bovine serum albumin (BSA; Solarbio, A8020) for 1 h at room temperature. Subsequently, cells were incubated with primary antibodies against Nanog (1:400; Rabbit; Proteintech, 14295-1-AP) and SSEA4 (1:400; Mouse; Abcam, ab16287) overnight at 4 °C.

Following three washes with PBS, cells were incubated with the corresponding secondary antibodies: Anti-mouse IgG (H + L), F(ab’)_2_ Fragment (Alexa Fluor 488 Conjugate; Cell Signaling Technology, 4408S) and Anti-rabbit IgG (H + L), F(ab’)_2_ Fragment (Alexa Fluor 555 Conjugate; Cell Signaling Technology, 4413S) at a dilution of 1:400 for 1 h at room temperature, protected from light. Nuclei were counterstained with Hoechst 33342 (Thermo Fisher Scientific, 62249). Finally, samples were mounted with fluorescence mounting medium (Servicebio, G1401) and stored at 4 °C until imaging. Images were acquired using an Olympus FV3000 confocal microscope.

### Generation of Embryoid Bodies (EBs)

To initiate differentiation, hiPSC colonies were dissociated into single cells. Briefly, cells were washed with PBS and incubated with 1 mL of Accutase (Biolegend, 423201) for 5 min, followed by neutralization and centrifugation at 200 × g for 3 min. Cells were resuspended and counted to achieve a final density of 5 × 10^4^ cells/mL in eTeSR medium supplemented with 10 μM Y-27632 dihydrochloride kinase inhibitor (MedChemExpress, HY-10583) and 50 ng/mL BMP4 (AcroBIOSYSTEMS, BM4-H5118). For EBs formation, 100 μL of the cell suspension was seeded into each well of a low-attachment 96-well U-bottom plate, yielding approximately 5,000 cells per well. The plates were centrifuged at 300g for 5 min to promote cell aggregation and incubated at 37 °C in 5% CO_2_ overnight to form EBs.

### Hematopoietic Differentiation

The differentiation process was carried out over a 15-day period. From day 0 to day 2, EBs were cultured in eTeSR (STEMCELL Technologies, 100-1215) supplemented with 50 ng/mL BMP4 (AcroBIOSYSTEMS, BM4-H5118) to induce mesoderm formation. On day 3, 12 EBs of the same genotype were transferred into each well of a low-attachment 12-well plate. Concurrently, the medium was replaced with a serum-free differentiation medium comprising StemPro-34 (Invitrogen, 10639011) supplemented with 2 mM L-Glutamine (Gibco, 35050061), 0.16 mM monothioglycerol (Sigma, M6145), 15.15 mg/mL holotransferrin (Sigma, T0665), 50 ng/mL BMP4 (AcroBIOSYSTEMS, BM4-H5118), 300 ng/mL Flt-3L (T&L Biotechnology, GMP-TL505-0050), 300 ng/mL SCF (T&L Biotechnology, GMP-TL504-0100), 10 ng/mL IL-3 (T&L Biotechnology, GMP-TL511-0050), and 10 ng/mL IL-6 (T&L Biotechnology, GMP-TL512-0100).

EBs were maintained in culture until day 15, with half-medium replenishment performed every 2–3 days (specifically on days 5, 8, 10, 12, and 14). This established differentiation protocol promotes the specification of mesodermal cells to bipotential hemato-endothelial progenitors (HEPs) between days 3 and 10. Subsequently, the protocol culminates in the commitment of HEPs to hematopoietic progenitor cells (HPCs; CD34^+^CD43^+^CD45^+^; days 10–15), myeloid progenitors (CD33^+^CD11b^+^), and monocytes (CD14^+^ within CD33^+^), recapitulating key milestones of early hematopoiesis.

### Declaration of Competing Interest

Z.L, Q.Z. and S.K. are inventors on a patent application (202411625725X) filed by Shenzhen Bay Laboratory that covers the design, composition and application of the split prime editing system and the fluorescence-guided enrichment strategy relating to this work. Other authors have no potential competing interests to declare.

### Data Availability

All data are available in the main text, supplementary information and **Source Data**. To facilitate broad accessibility, the key plasmids generated in this work have been deposited in the Addgene (https://www.addgene.org/) and WeKwikGene (https://wekwikgene.wllsb.edu.cn/labs/a0055f03-cbf9-4d60-8e0b-834140041f57) repositories. Additional information related to this study is available from the corresponding author upon reasonable request.

## Acknowledgements

This work was supported by the Shenzhen Bay Laboratory Startup Fund and Major Project Grant (S241101007-2); Postdoctoral research funding provided by the Human Resources and Social Security Bureau of Shenzhen Municipality; the Spanish Ministry of Science and Innovation (Ministerio de Ciencia e Innovación) / State Research Agency (AEI) through the Retos Investigación grant (PID2023-146290OB-I00) and the “Programa FORTALECE” (FORT23/00032); and the Carlos III Health Institute (ISCIII) through AES 2023 (Award no. AC23_2/00040) within the European Joint Programme on Rare Diseases framework. We thank the members of the Z.L. and A.G. laboratories for insightful discussions. We are grateful to the Research Core Facility at Shenzhen Bay Laboratory for technical assistance, and to the laboratories of Lin Deng and Joe Z. Zhang for generously sharing reagents and protocols. We apologize for not being able to cite additional work owing to space limitations.

## Author Contributions

**Zhichao Qiu**: Methodology, Investigation, Data Curation, Formal Analysis, Visualization, Writing – Original Draft, Writing – Review & Editing. **Keke Sun**: Methodology, Investigation, Data Curation, Formal Analysis, Visualization. **Qingwei Zeng**: Investigation, Data Curation. **Ziwei Luo**: Methodology, Data Curation, Formal Analysis. **Xinran Liu**: Investigation, Data Curation. **Yaping Li**: Investigation, Data Curation, Formal Analysis. **Xiang Lei**: Methodology, Investigation, Data Curation. **Ruilin Zhao**: Methodology, Investigation, Data Curation. **Zhen Zhang**: Methodology, Investigation, Data Curation. **Damia Romero-Moya**: Methodology, Investigation, Data Curation, Formal Analysis. **Yuan Yang**: Investigation, Data Curation. **Xuan Wang**: Methodology, Data Curation. **Ehsan Hashemi**: Investigation. **Joe Z. Zhang**: Resources, Supervision. **Xiaolong Wang**: Resources, Supervision. **Alessandra Giorgetti**: Investigation, Methodology, Resources, Supervision, Writing – Review & Editing, Funding acquisition. **Zhuobin Liang**: Conceptualization, Investigation, Methodology, Supervision, Writing – Original Draft, Writing − review & editing, Project administration, Funding acquisition.

## Extended Data

Source Data

## Supplementary Information

Supplementary Tables 1-5

**Supplementary Figure 1.**
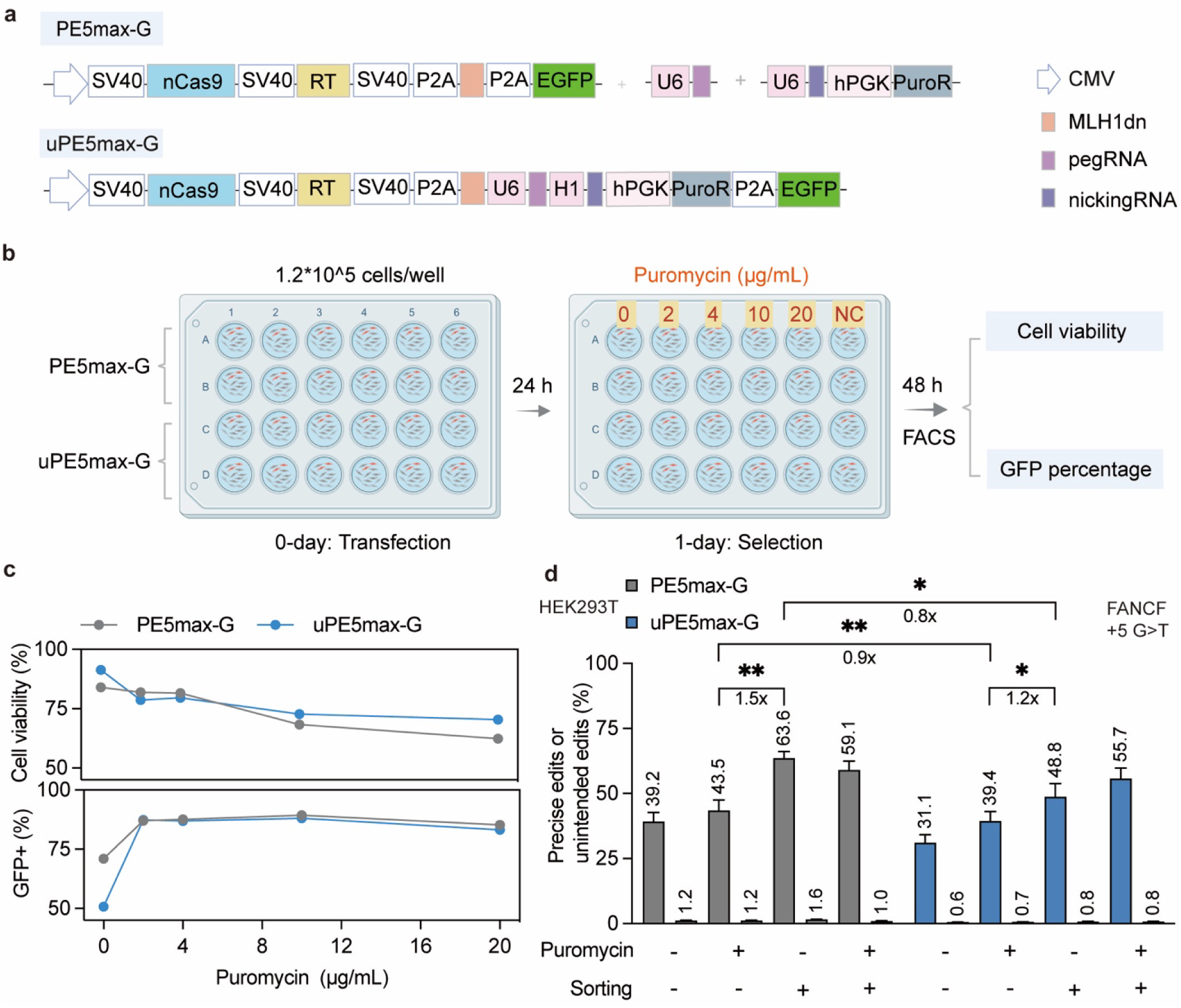
Puromycin selection yields incomplete enrichment and high cytotoxicity during prime editing. **a**, Schematics of the PE5max-G and uPE5max-G vector architectures. PE5max-G incorporates a P2A-linked EGFP reporter, while uPE5max-G consolidates all protein and RNA components into a single vector. **b**, **c**, Experimental workflow (**b**) and flow cytometric quantification (**c**) of cell viability and transfection efficiency (GFP+ %) across varying puromycin concentrations. **d**, Frequencies of precise prime edits (large bars) and unintended edits (small bars) induced by PE5max-G (gray) and uPE5max-G (blue) at the FANCF +5 G>T locus in HEK293T cells. Cells were subjected to no enrichment, Puromycin selection (+), FACS sorting (+), or a combination of both (+/+). Data are presented as mean ± s.e.m. (n = 2 independent biological replicates for **c**; n = 3 independent biological replicates for **d**). Fold changes between indicated groups are shown below the brackets. Statistical significance was determined by paired two-tailed Student’s t-test (* p < 0.05, ** p < 0.01).

**Supplementary Figure 2.**
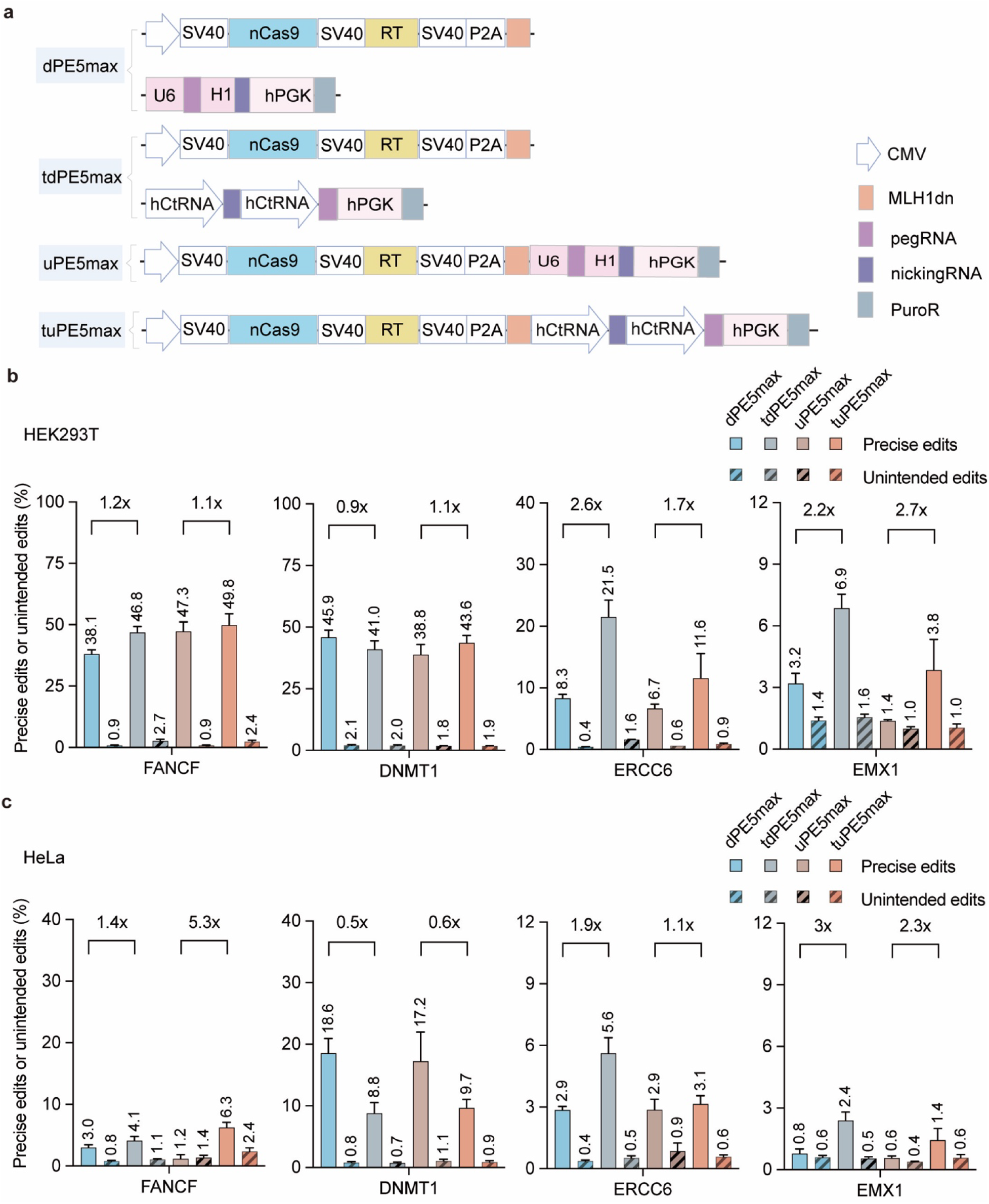
Validation of hCtRNA-driven RNA arrays in dual and unified vector architectures. **a**, Schematics of dual-vector (dPE5max, tdPE5max) and unified (uPE5max, tuPE5max) prime editing systems. The tdPE5max and tuPE5max variants utilize hCtRNA promoters to drive the RNA elements. **b**, **c**, Editing efficiencies and unintended edit frequencies across four genomic loci (*FANCF, DNMT1, ERCC6, EMX1*) in HEK293T (**b**) and HeLa (**c**) cells. Fold changes compare the hCtRNA-driven systems to their U6/H1-driven counterparts. Editing outcomes were quantified by targeted deep sequencing. Data are presented as mean ± s.e.m. (n = 3 independent biological replicates) .

**Supplementary Figure 3.**
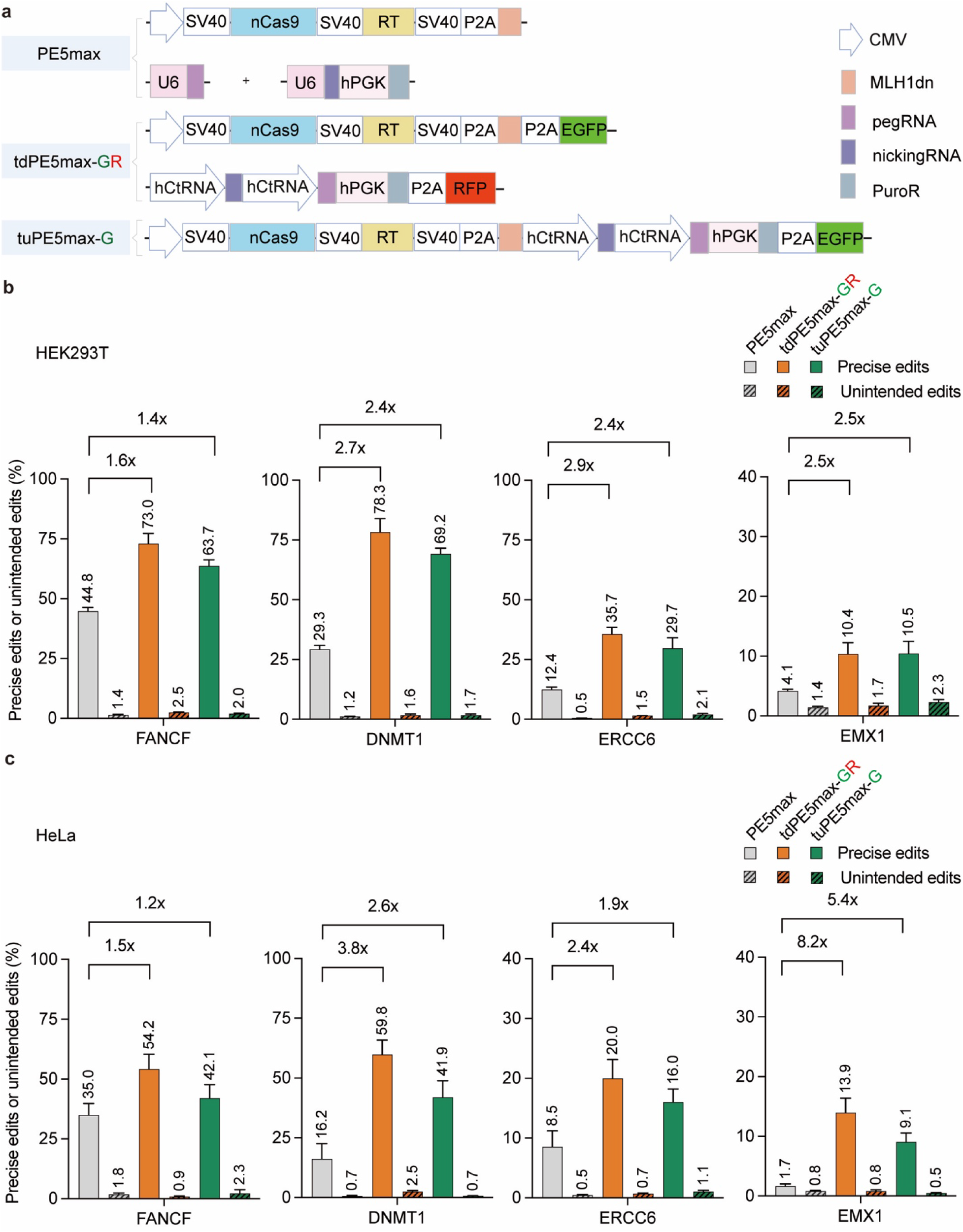
Fluorescence-based enrichment improves editing yields independent of vector topology. **a**, Schematics of PE5max, tdPE5max-GR, and tuPE5max-G architectures. The GR and G variants incorporate P2A-linked fluorescent reporters for sorting. **b**, **c**, Prime editing efficiencies and unintended edit frequencies at four loci in HEK293T (**b**) and HeLa (**c**) cells. Fluorescent cell populations were sorted by FACS at 48 h post-transfection. Fold improvements relative to the standard PE5max control are indicated above the bars. Editing outcomes were quantified by targeted deep sequencing. Data are presented as mean ± s.e.m. (n = 3 independent biological replicates).

**Supplementary Figure 4.**
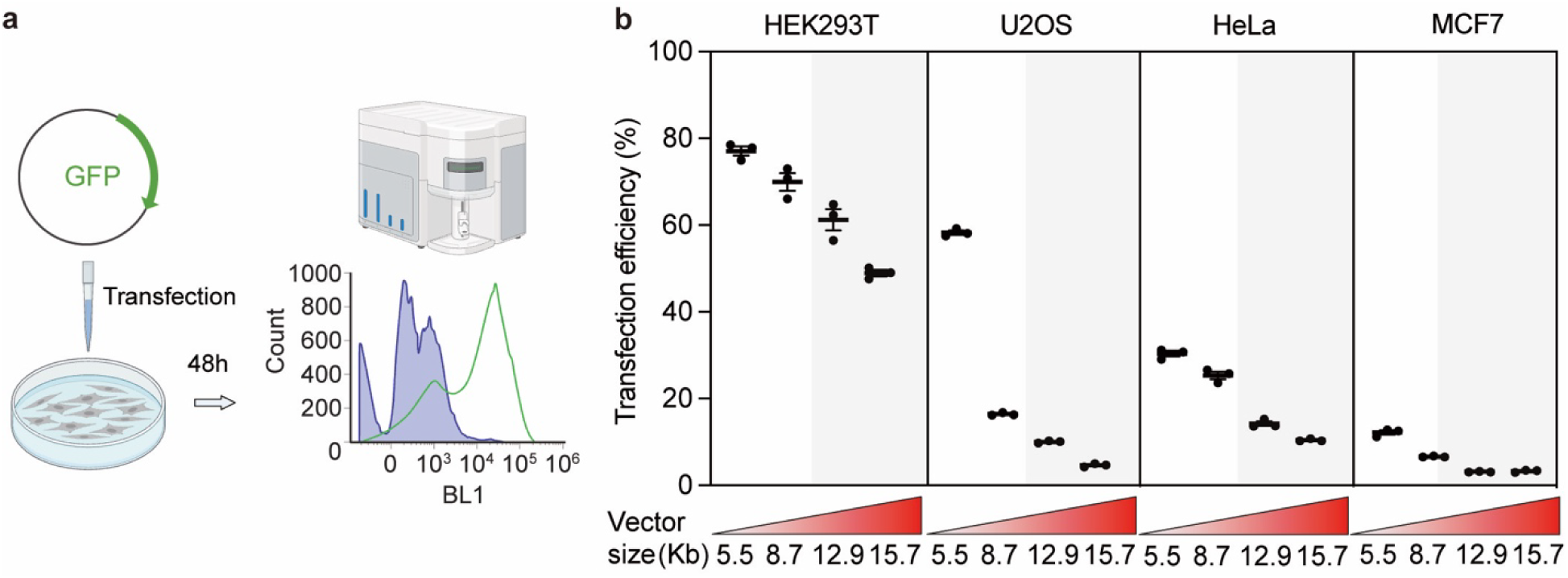
Plasmid size inversely correlates with transfection efficiency in mammalian cell lines. **a**, Experimental workflow for assessing transfection efficiency via flow cytometry at 48 h post-transfection. **b**, Transfection efficiencies of GFP-encoding plasmids of varying sizes (5.5, 8.7, 12.9, and 15.7 kb) across HEK293T, U2OS, HeLa, and MCF7 cells. Gray shading highlights regions of diminished transfection efficiency corresponding to larger vector payloads. Data are presented as mean ± s.e.m. (n = 3 independent biological replicates) .

**Supplementary Figure 5.**
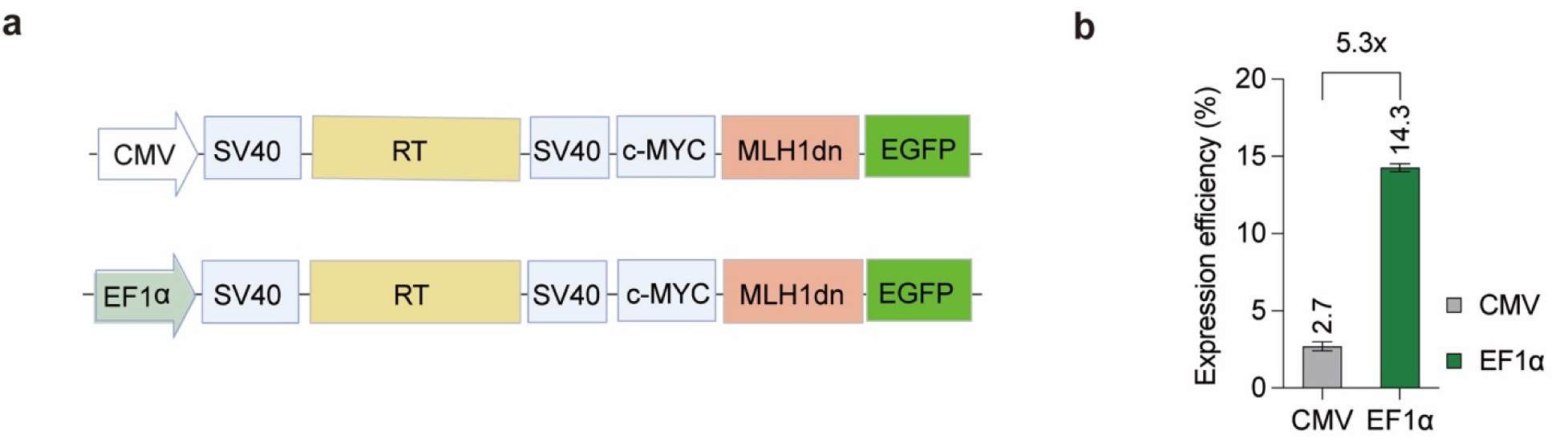
EF1α promoter drives superior transgene expression compared to CMV promoter in hiPSCs. **a**, Schematics of expression vectors utilizing either the CMV or EF1α promoter to drive an EGFP-tagged reporter. **b**, Flow cytometric quantification of EGFP-positive hiPSCs at 48 h post-transfection. The EF1α promoter demonstrated a 5.3-fold increase in expression efficiency over the CMV promoter. Data are presented as mean ± s.e.m. (n = 3 independent biological replicates) .

**Supplementary Figure 6.**
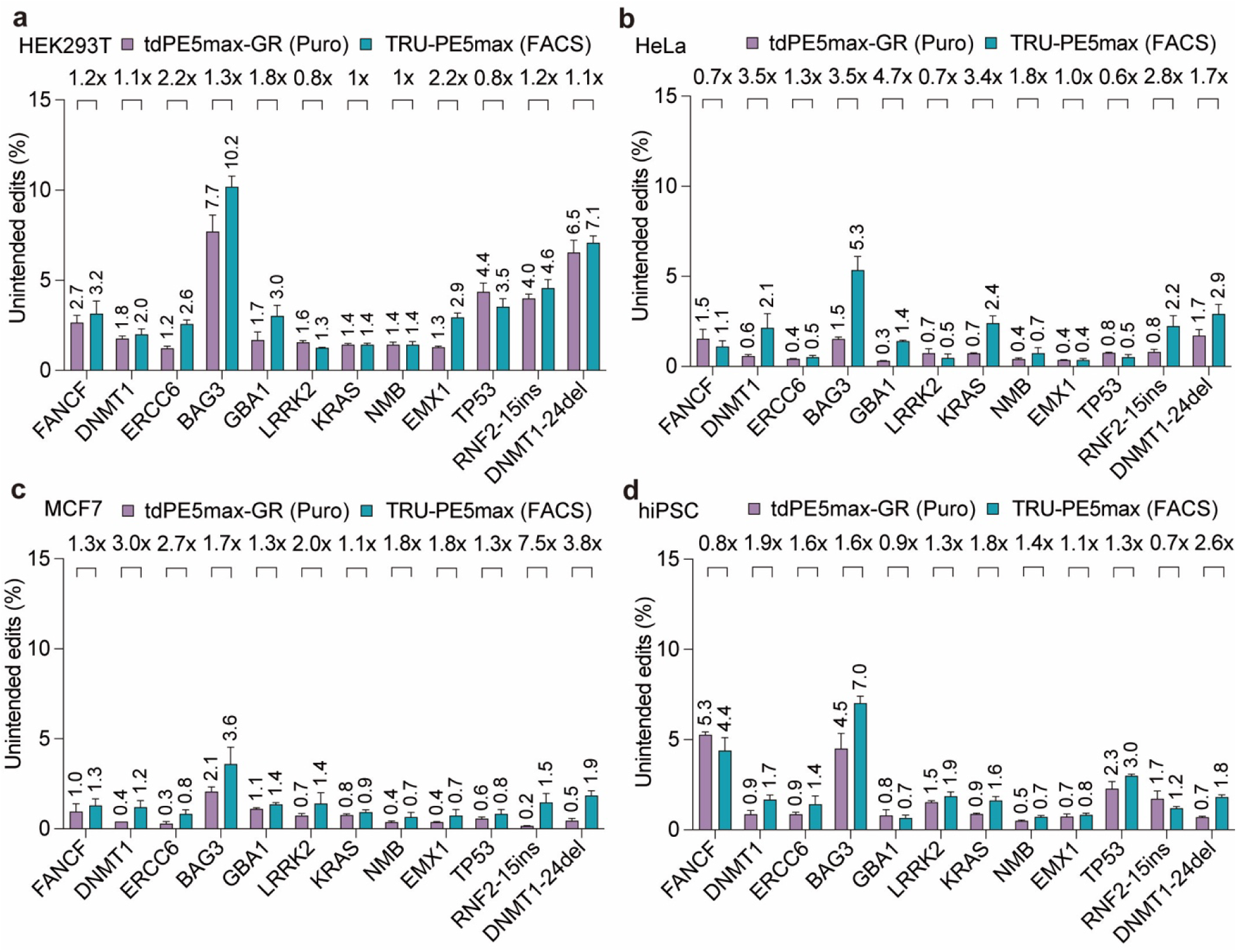
Unintended edit profiles of TRU-PE5max versus puromycin-selected controls (related to Fig. 1b–e). **a**–**d**, Unintended edit frequencies at 12 genomic loci following treatment with tdPE5max-GR (puromycin selection) or the TRU-PE5max (FACS enrichment) system in HEK293T (**a**), HeLa (**b**), MCF7 (**c**), and hiPSCs (**d**). Puromycin was applied at 4 μg/mL for cancer lines and 0.5 μg/mL for hiPSCs. Fold changes between systems are indicated above the bars. Data are presented as mean ± s.e.m. (n = 3 independent biological replicates) .

**Supplementary Figure 7.**
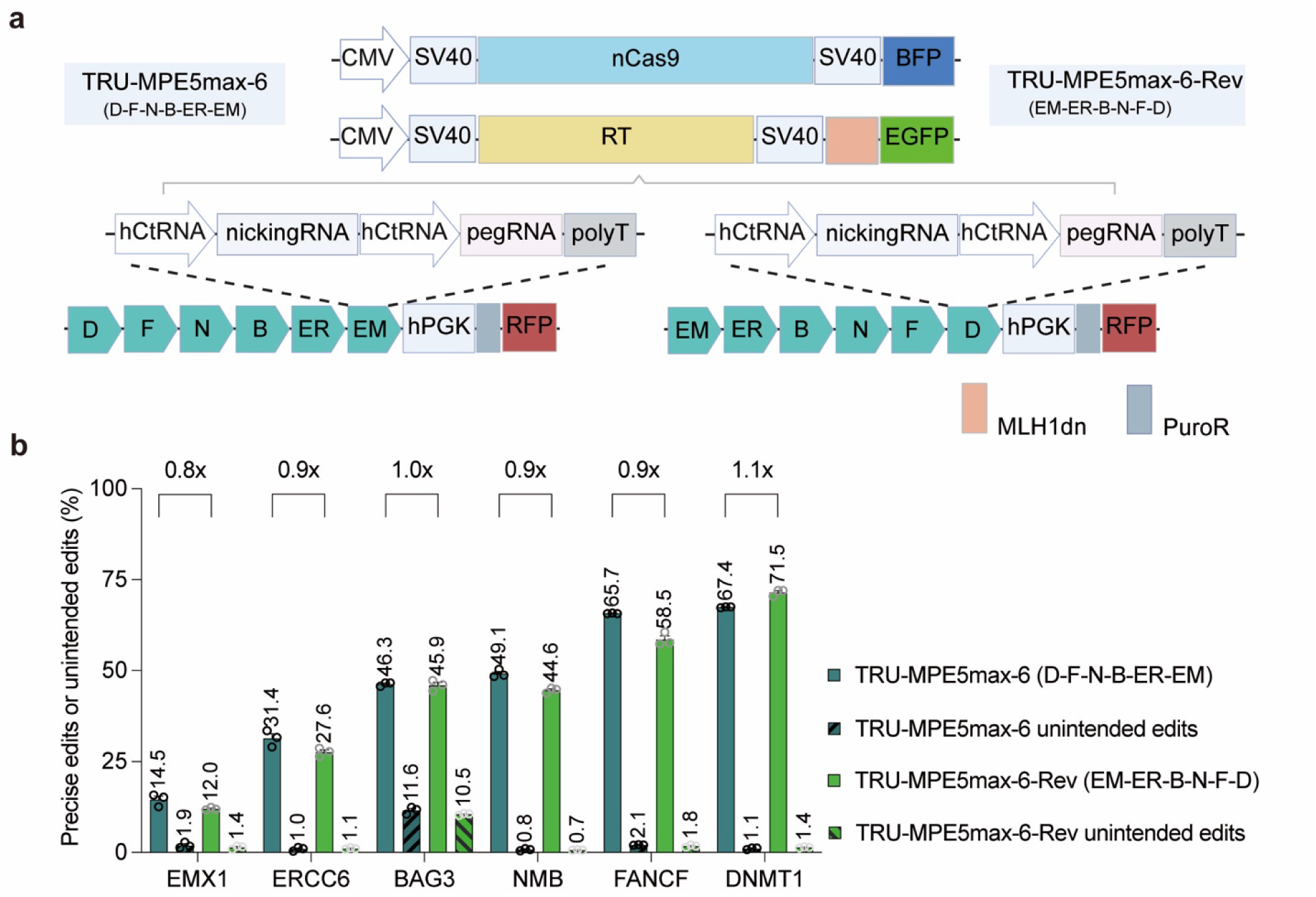
Impact of pegRNA position within hCtRNA arrays on TRU-PE multiplex editing efficiency. **a**, Schematics of the TRU-MPE5max-6 vectors featuring two distinct pegRNA arrangements. Target genes were organized in either a forward (D-F-N-B-ER-EM) or reverse (EM-ER-B-N-F-D) sequence within the hCtRNA-driven array (D: *DNMT1*; F: *FANCF*; N: *NMB*; B: *BAG3*; ER: *ERCC6*; EM: *EMX1*). **b**, Comparative analysis of editing efficiencies and unintended edit frequencies between the forward and reverse orientations in HEK293T cells. Fold changes relative to the forward order are indicated. Data are presented as mean ± s.e.m. (n = 3 independent biological replicates) .

**Supplementary Figure 8.**
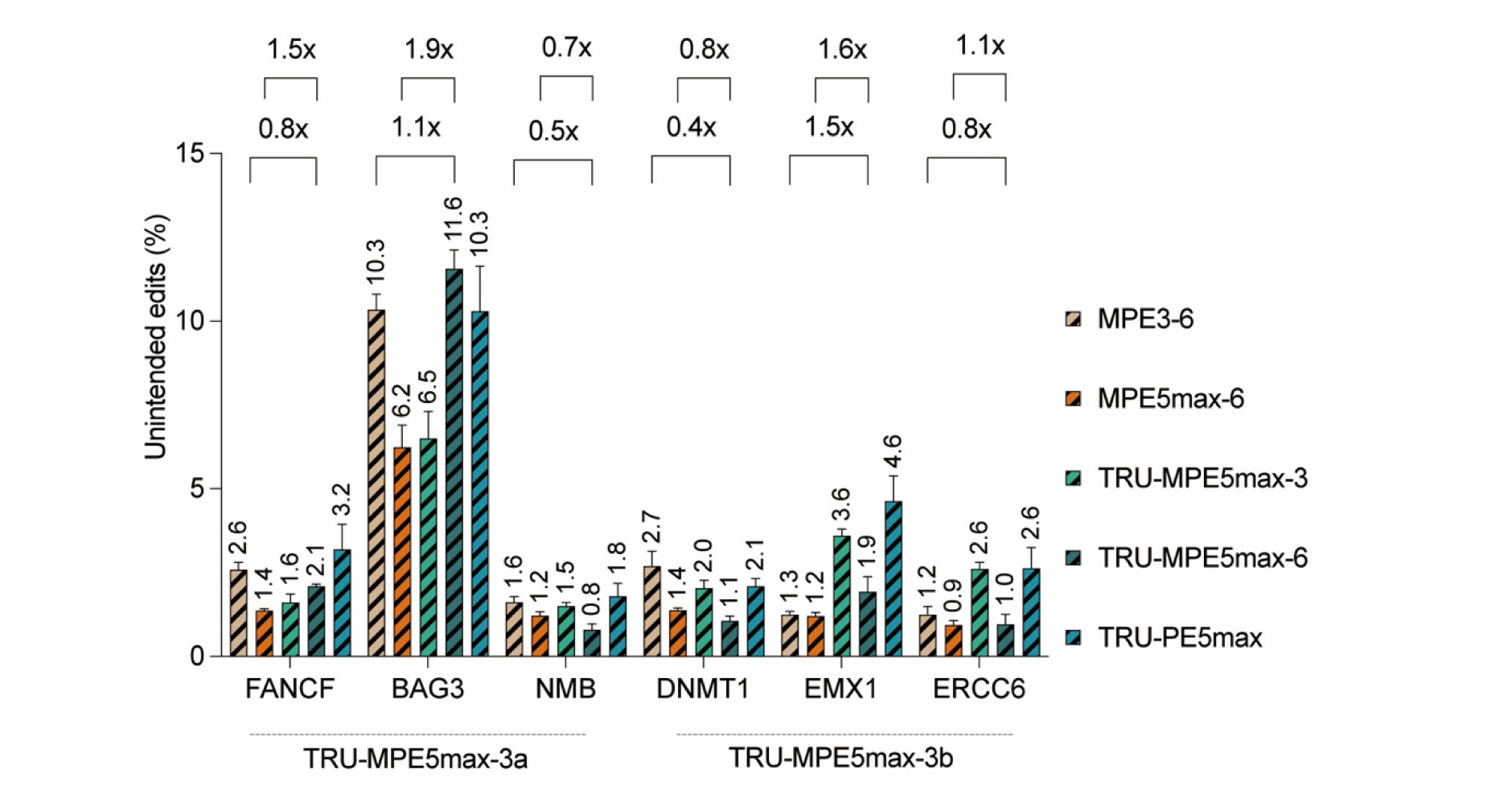
Analysis of unintended byproduct frequencies during multiplex prime editing (related to Fig. 2b). Comparison of unintended edit frequencies across six genomic loci in HEK293T cells using various multiple editing configurations. Systems evaluated include standard multiplex controls (MPE3-6, MPE5max-6), alongside the TRU-MPE5max (3-plex arrays a and b, and 6-plex) and the single-plex controls (TRU-PE5max). Fold changes relative to controls are shown above the bars. Data are presented as mean ± s.e.m. (n = 3 independent biological replicates) .

**Supplementary Figure 9.**
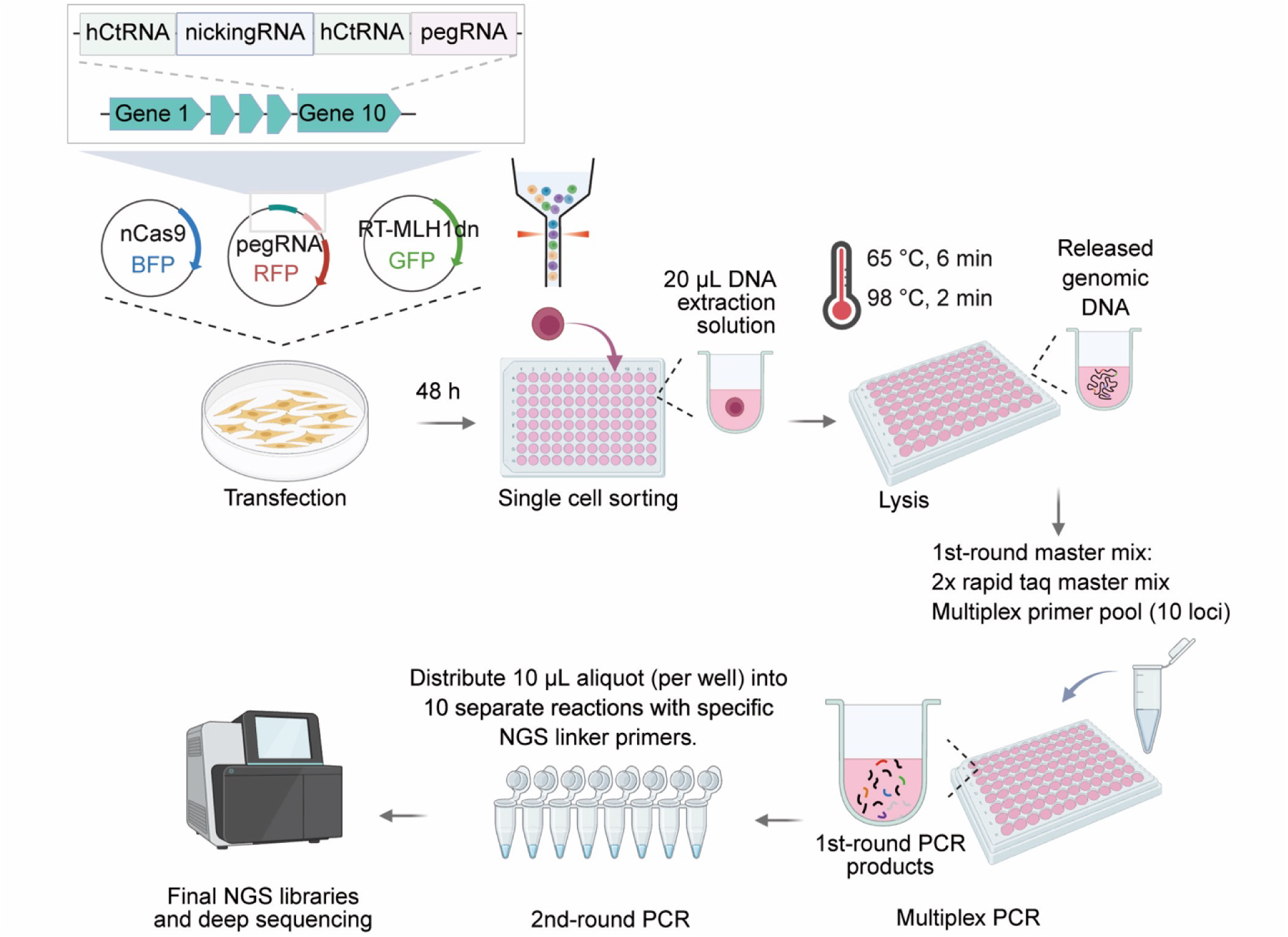
Workflow for single-cell targeted amplicon sequencing of multiplexed loci (related to Fig. 2d–f). Schematic of the streamlined pipeline to quantify multiplex editing efficiencies at single-cell resolution. At 48 h post-transfection with TRU-MPE5max components, triple-fluorescence-positive (BFP+/GFP+/RFP+) single cells are sorted via FACS into 96-well plates containing DNA extraction solution. Following thermal lysis, genomic DNA undergoes a first-round multiplex PCR amplification, followed by a second-round PCR to incorporate specific NGS linker primers before pooling for targeted deep sequencing.

**Supplementary Figure 10.**
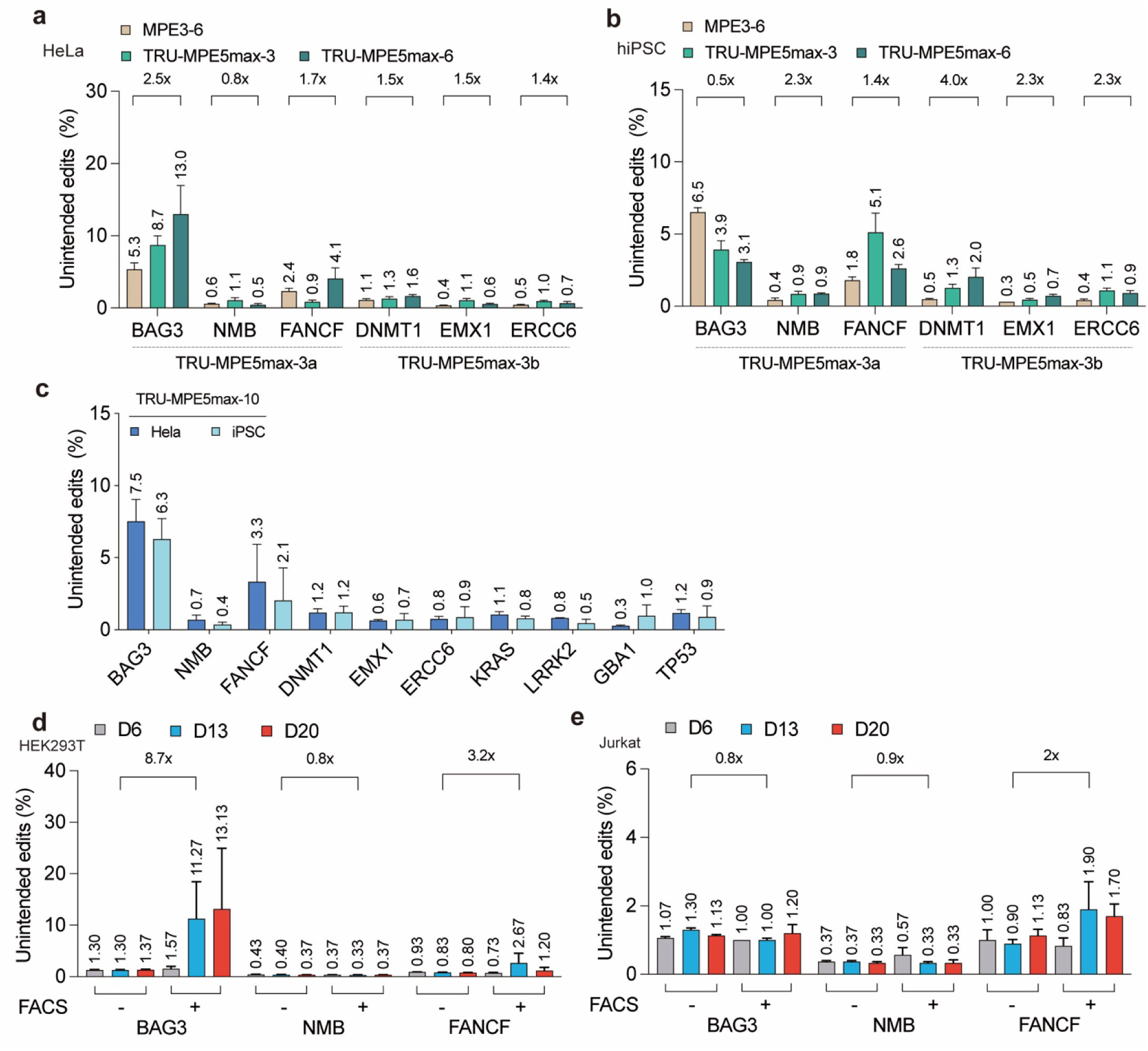
Unintended edit profiles of high-order multiplex editing in HeLa and hiPSC lines (related to Fig. 3a–e). **a**, **b**, Unintended edit frequencies at six genomic loci utilizing two distinct 3-plex arrays (TRU-MPE5max-3a/b) and a 6-plex array in HeLa (**a**) and hiPSCs (**b**). **c**, Unintended edit frequencies following simultaneous 10-locus editing (TRU-MPE5max-10) in HeLa and hiPSC cells. Data are presented as mean ± s.e.m. (n = 3 independent biological replicates). **d**, **e**, Unintended edit frequencies of lentiviral delivery for 3-plex unintended edits (*BAG3*, *NMB*, *FANCF*) in HEK293T (**d**) and Jurkat T cells (**e**).

**Supplementary Figure 11.**
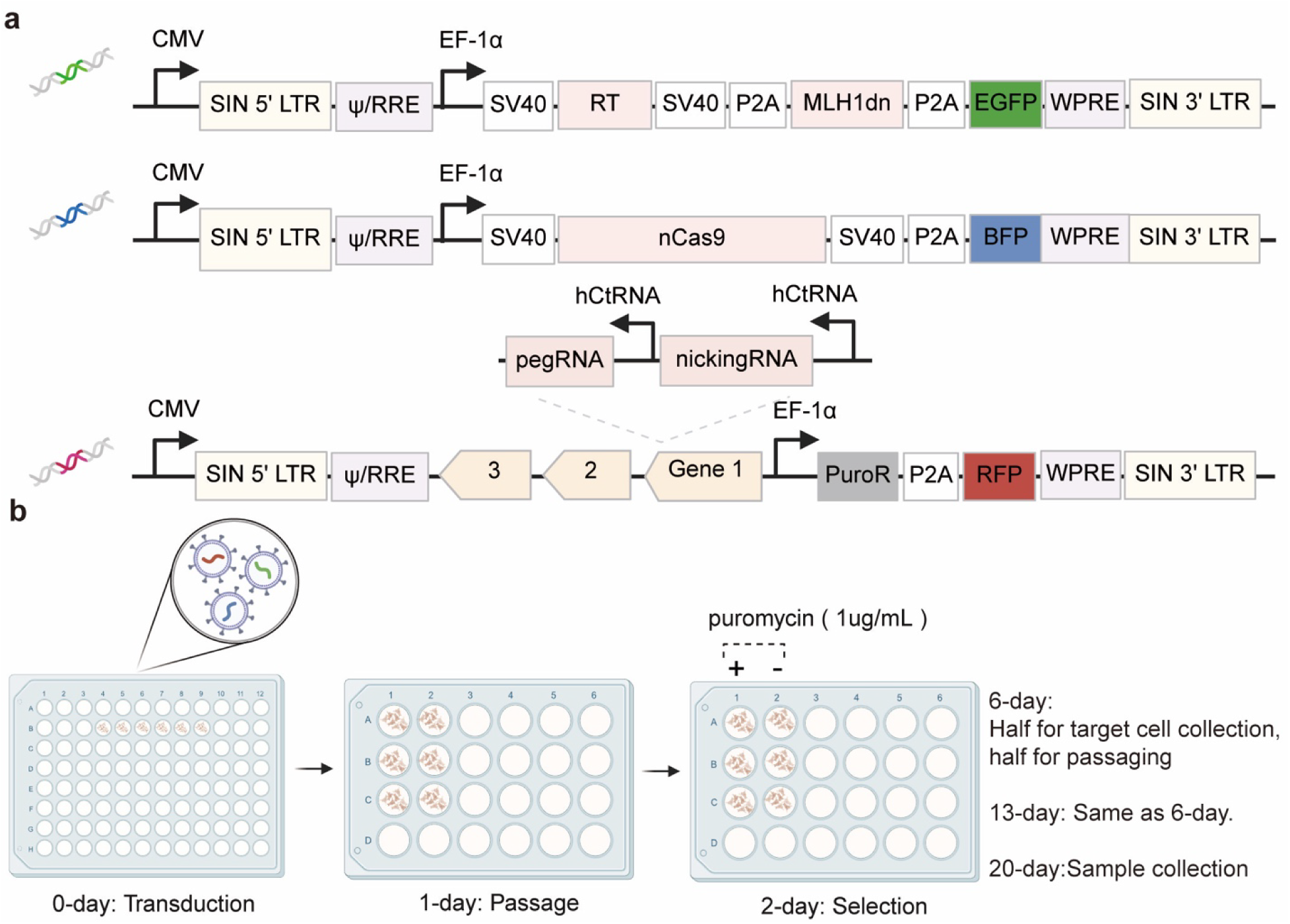
Viral vector design and transduction workflow for TRU-MPE delivery (related to Fig. 3d,e). **a**, Schematics of lentiviral transfer plasmids for the tripartite TRU-MPE5max-3 system. The vectors independently encode RT-MLH1dn (EGFP) and nCas9 (BFP) under EF-1α promoters, while the 3-plex hCtRNA-driven array vector (RFP) co-expresses a puromycin resistance cassette. **b**, Experimental timeline indicating viral transduction (day 0), passage (day 1), initiation of puromycin selection (1 μg/mL, day 2), and subsequent sample collection at days 6, 13, and 20 post-transduction.

**Supplementary Figure 12.**
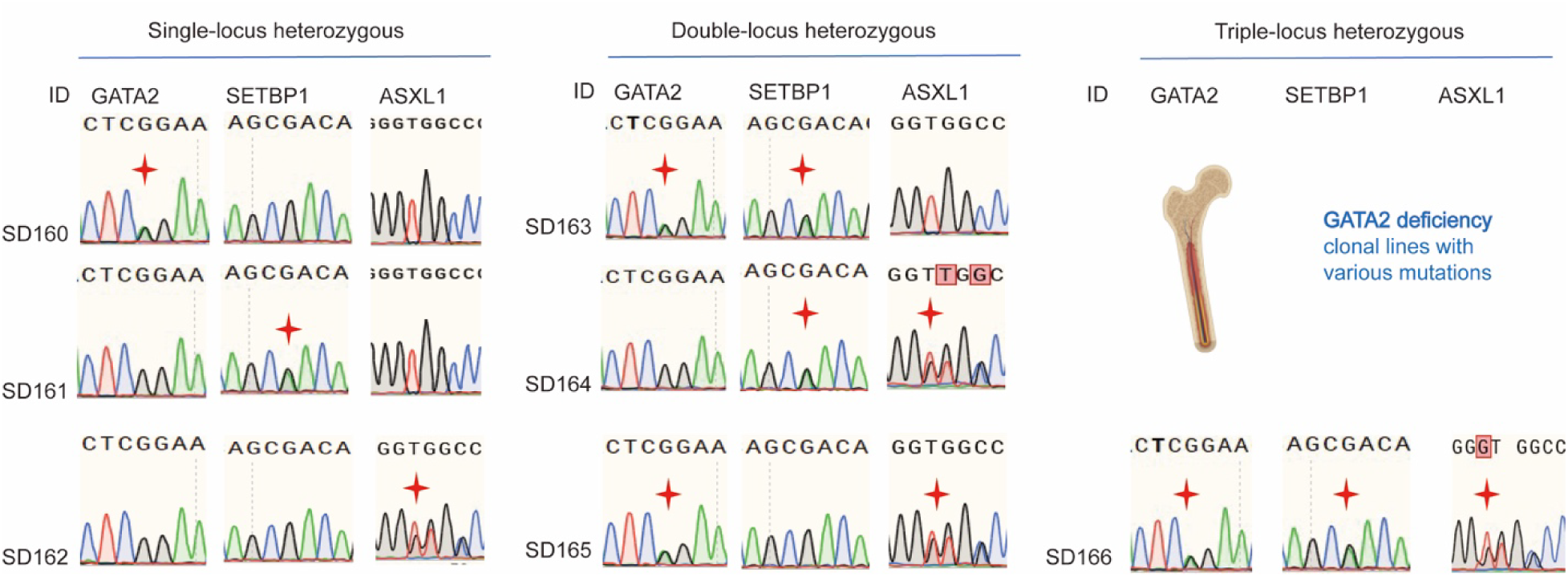
Genotypic validation of the isogenic GATA2-deficiency hiPSC lines (related to Fig. 6c). Representative Sanger sequencing chromatograms confirming modifications at the *GATA2*, *SETBP1*, and *ASXL1* loci within isolated hiPSC clones (SD160–SD166). Red stars mark the intended mutations, validating the successful generation of single-locus (SD160–SD162), double-locus (SD163–SD165), and triple-locus (SD166) heterozygous edits.

## Notes

### Competing Interest Statement

Z.L, Z.Q. and K.S. are inventors on a patent application (202411625725X) filed by Shenzhen Bay Laboratory that covers the design, composition and application of the split prime editing system and the fluorescence-guided enrichment strategy relating to this work. Other authors have no potential competing interests to declare.

